# CIS calibrates GM-CSF signalling strength to regulate macrophage polarization via a STAT5-IRF8 axis

**DOI:** 10.1101/2022.03.31.486495

**Authors:** Shengbo Zhang, Jai Rautela, Naiara G Bediaga, Tatiana B Kolesnik, Yue You, Laura F Dagley, Justin Bedo, Hanqing Wang, Li Sun, Robyn Sutherland, Elliot Surgenor, Nadia Iannarella, Rhys Allan, Fernando Souza-Fonseca-Guimaraes, Yi Xie, Qike Wang, Yuxia Zhang, Yuekang Xu, Stephen L Nutt, Andrew M Lew, Nicholas D Huntington, Sandra E Nicholson, Michaël Chopin, Yifan Zhan

## Abstract

The cytokine granulocyte-macrophage-colony stimulating factor (GM-CSF) possesses the ability to differentiate macrophages (MØs) with opposing functions, namely proinflammatory M1-like and immunosuppressive M2-like. Despite the importance and opposing functional outcomes of these processes, the intrinsic mechanism that regulates the functional polarization of MØs under GM-CSF signaling remains elusive. Here we show that GM-CSF induced MØs polarisation resulted in the expression of the Cytokine-inducible SH2-containing protein (CIS), and that CIS deficiency diverted differentiation of monocytes into immunosuppressive M2-like MØs expression. CIS deficiency resulted in the hyperactivation of the JAK-STAT5 signaling pathway, consequently promoting the downregulation of the transcription factor Interferon Regulatory Factor 8 (IRF8). Loss and gain of function approaches highlighted IRF8 as a critical instructor of the M1-like polarisation program. In vivo, CIS deficiency led to skewing to M2-like macrophages, which induced strong Th2 immune responses characterised by the development of severe experimental asthma. Collectively, we reveal a CIS-censored mechanism interpreting the opposing actions of GM-CSF in MØ differentiation and uncovering its role in controlling allergic inflammation.

## Introduction

Macrophages (MØs) are key immune cells that play crucial role in immune defence against invading pathogens, but also have critical functions in regulating and maintaining tissue homeostasis(Lavin et al., 2015). It has long been appreciated that MØs show functional plasticity/polarization and dynamically respond to different physiological situations. External cues, including cytokines and Toll-like receptor (TLR) agonists can facilitate MØs functional polarization(Murray et al., 2014). Historically, interferon (IFN)-γ or IL-4 had been used respectively to polarize classically activated M1 MØs with strong proinflammatory functions or alternatively activated M2 MØs with anti-inflammatory functions (Murray et al., 2014) .

Differing from the aforementioned modes of MØ polarization, MØs generated from bone marrow progenitors or monocytes in presence of granulocyte-macrophage-colony stimulating factor (GM-CSF) or macrophage-colony stimulating factor (M-CSF) feature both M1 and M2 like characteristics, respectively (Fleetwood et al., 2007). GM-CSF differentiated MØs are thought to be typically M1-like, producing pro-inflammatory cytokines upon stimulation with TLR ligands(Fleetwood et al., 2007; Martinez and Gordon, 2014; Verreck et al., 2004). Corroborating evidence in autoimmune disease supports a pro-inflammatory role for GM-CSF in vivo (reviewed in(Becher et al., 2016; Hamilton, 2008)). Adding to its pro-inflammatory properties, GM-CSF has long been used for its strong immune adjuvant attributes in a vaccination setting in order to promote anti-tumor immunity (Dranoff et al., 1993).Paradoxically, GM-CSF has also been associated with the development of suppressive M2-like MØs in various tumour settings (Bayne et al., 2012; Bronte et al., 1999; Sielska et al., 2013) and after renal ischemia (Huen et al., 2015). GM-CSF also has a critical role in the induction of allergic inflammation (Cates et al., 2004; Willart et al., 2012), and consequently its neutralization dampened inflammation in certain allergic settings (Cates et al., 2004), although it is not clear from these studies whether M2-like MØs are directly responsible for the induction of allergic inflammation. Nevertheless, M2-like MØs are known for their ability to support Th2 immunity (Mills et al., 2000). In murine models of dextran sulfate sodium (DSS)-induced colitis, a protective role of GM-CSF was also associated with GM-CSF induced myeloid cells(Bernasconi et al., 2010). Together, these studies point to the paradoxical functional outcomes driven by GM-CSF signalling in MØs, which can result in many pathological settings, yet the molecular mechanism governing the functional dichotomy of GM-CSF action on MØs is largely unknown.

GM-CSF signaling strength and duration is tighly regulated by the induction of suppressors of cytokine signaling (SOCS) proteins which limit cytokine induced signaling (Alexander and Hilton, 2004). Among the SOCS family members, cytokine-inducible SH2 protein (CIS), is a known target of STAT5 activation(Feldman et al., 1997; Matsumoto et al., 1997) which is induced by GM-CSF in MØs (Feldman et al., 1997; Lehtonen et al., 2002). CIS deficient mice housed in a specifc pathogen-free environment show no overt defects in myelopoiesis, suggestive of low GM-CSF or CIS expression in unchallenged situations (Marine et al., 1999; Matsumoto et al., 1999). In line with this, the impact of CIS deficiency in NK or T cells only became apparent following stimulation with exogenous IL-15 or TCR engagement, respectively (Delconte et al., 2016; Palmer et al., 2015; Yang et al., 2011). Thus, a role for CIS in regulating macrophages polarisation under inflammatory settings remained largely unexplored.

Here we explored the role of CIS in regulating MØ functional polarization following GM-CSF stimulation. We found that CIS deficiency biased macrophages toward M2-like MØs, with strong immunosuppressive functions, limited IL-12 production and Th2 bias. The functional diversion was largely the result of the hyperactivation of the JAK-STAT5 signaling pathway and reduced IRF8 induction. In vivo *Cish^−/−^* MØs primed a strong Th2 response and promoted excess inflammation in allergic asthma independent from the previsouly described T-cell intrinsic role of CIS in regulation of Th2 immunity (Yang et al., 2013). Thus, our study highlights a critical role for CIS in fine tuning GM-CSF signaling and in promoting/ maintaining M1 like features in macrophages.

## Results

### CIS deficiency leads to the generation of MØs that strongly inhibit T-cell responses

In line with previous studies (Marine et al., 1999; Palmer et al., 2015), we did not observe any conspicuous defects in multiple types of myeloid cell sub-types or BM hematopoietic progenitors in *Cish^−/−^* mice (**Figure S1&S2**). To investigate the role of CIS in MØ differentiation, we cultured BM progenitors from WT and *Cish*^−/−^ mice (Palmer et al., 2015) for 7 days with GM-CSF to generate MØs (GM-MØs; CD11c^+^MHCII^int^ CD11b^hi^CD115^hi^CD86^lo^Flt3^lo^) (up to 90% of CD11c^+^ cells) and DCs (GM-DCs; CD11c^+^MHCII^hi^ CD11b^int^CD115^−^CD86^hi^Flt3^hi^) (Helft et al., 2015) (**Figure 1A**). GM-MØs were dominant for both genotypes (80-90% of total CD11c^+^ cells), with a small but significant reduction in the proportion of GM-DCs in *Cish*^−/−^ BM cell cultures (**Figure 1A**). Flow cytometry and fluorescence microscopy revealed that *Cish*^−/−^ GM-MØs were larger in size than WT GM-MØs (**Figure 1B).** We also found that *Cish*^−/−^ GM-MØs had lower cell surface abundance of CD115 (M-CSF receptor; M-CSFR) compared to WT GM-MØs, while GM-MØs of both genotypes expressed similar levels of GM-CSFR α and β (**Figure 1C**). Downregulation of CD115 by GM-CSF was dose-dependent and cell-intrinsic since *Cish*^−/−^ GM-MØs derived from co-culture with WT cells also had lower CD115 expression (**Figure S3A&B**). Lower expression of other myeloid markers such as CD209, CCR2, F4/80 and CD14 by *Cish*^−/−^ GM-MØs were also observed (**Figure S3B&C**), while GM-DCs of both genotypes had similar expression of multiple cell surface markers (**Figure S3D**).

**Figure 1.**
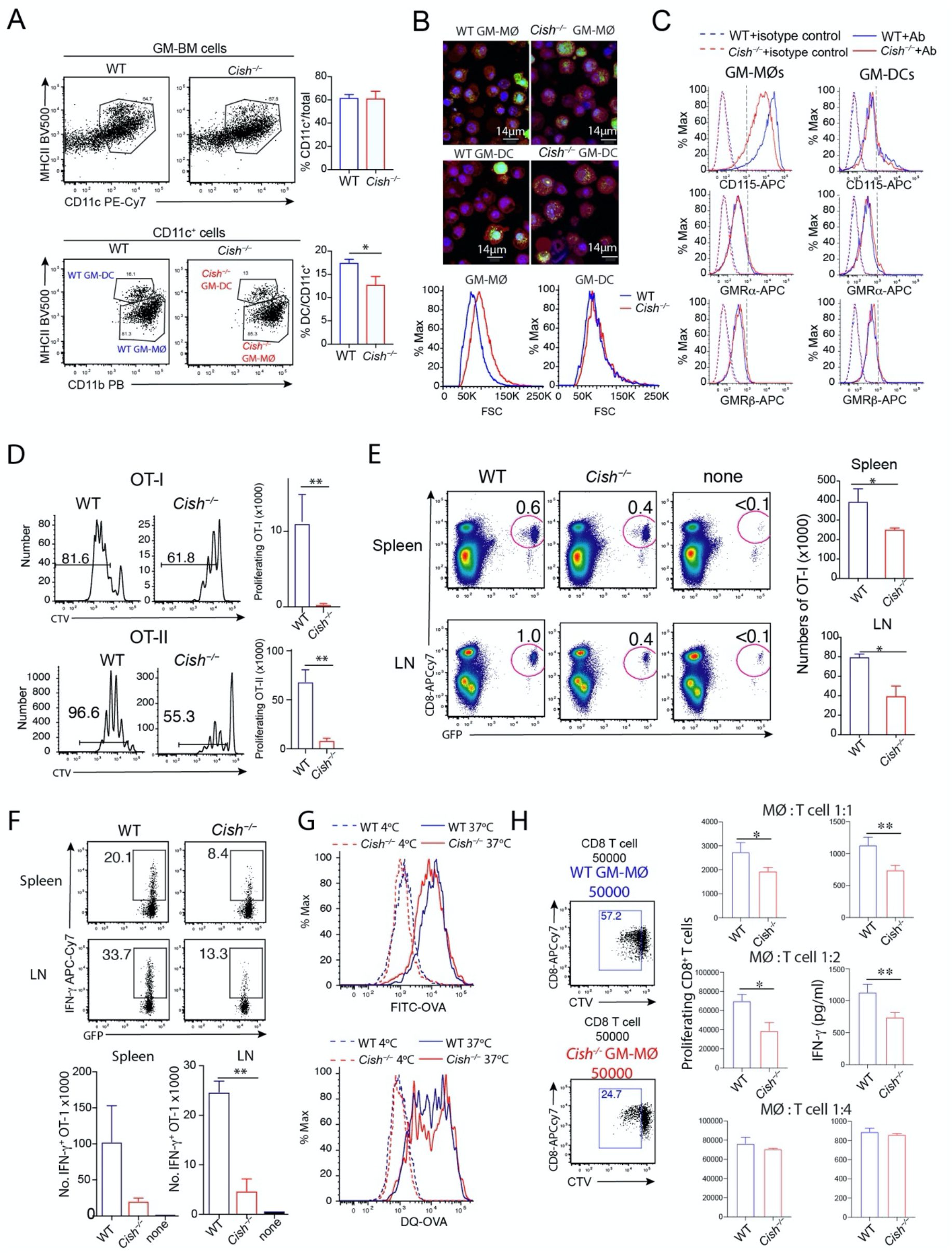
*Cish*^−/−^ GM-MØs are less capable of inducing T cell responses. (**A**) WT and *Cish*^−/−^ GM-MØs BM cells were cultured with 10ng/mL GM-CSF for 7 days. Harvested cells were analysed for GM-MØs and GM-DCs. (**B**) Sorted GM-MØs and GM-DCs were stained for CellTracker velvet (Red), LysoTracker green (green) and SIR-DNA (blue). Histograms show cell size by forward scatter of WT and *Cish*^−/−^ GM-MØs and GM-DCs. (**C**) Histograms show expression of M-CSF (CD115) and GM-CSF receptors by WT and *Cish*^−/−^ GM-MØs and GM-DCs. (**D**) CTV labelled OT-I T cells or OT-II were cultured with WT or *Cish*^−/−^ GM-MØs in presence of ovalbumin (OVA). Bar graphs represent the mean numbers ± SD of proliferating OT-I T cells at 48h (right) and OT-II T cell proliferation 60h. Data are representative of three independent experiments. (**E**) B6 mice that had been previously injected with GFP-OT-I cells, were intravenously infused with OVA-pulsed WT (n=3) or *Cish*^−/−^ GM-MØs (n=3). *P<0.05, by t test. Antigen-induced T cell expansion was evaluated 5 days after MØ transfer. (**F**) IFNγ production by GFP-OT-I was evaluated after 4 hr stimulation with PMA/Ionomycin. **P<0.01 by t test. Data are representative of two independent experiments. (**G**) Uptake and processing soluble OVA by WT and *Cish*^−/−^ GM-MØs. For uptake, cells were incubated with FITC‐OVA at 37°C or kept on ice for 30 min. For processing, cells were incubated with DQ-OVA at 37°C for 30 min. Then samples were either incubated at 37°C for further 90 min or kept on ice. (**H**) Purified CD8^+^ T cells from B6 mice were stimulated with anti-CD3/anti-CD28 with indicated number of WT or *Cish*^−/−^ GM-MØs for 3 days. Cell proliferation and cytokine production were then determined. *P<0.05, **P<0.01 by t test.

Next, we sought to compare the functions of GM-MØs generated from WT or *Cish*^−/−^ mice. Firstly, we evaluated GM-MØs for their ability to stimulate antigen-specific T cell proliferation. GM-MØs were pulsed with Ovalbumin (OVA) and subsequently co-cultured with dye-labelled MHCI and MHCII restricted OVA-specific CD8^+^ or CD4^+^ T cells (OT-I and OT-II, respectively). We observed that T-cell proliferation was significantly weaker when T cells were co-cultured with OVA-pulsed *Cish*^−/−^ GM-MØs compared to T cells co-cultured with OVA-pulsed WT GM-MØs (**Figure 1D, Figure S3E**). A similar observation was made for GM-DCs **(Figure S3E**), suggesting that CIS expression in MØs is necessary to promote T cell expansion. In line with the above, adoptive transfer of OVA-pulsed GM-MØs into C57BL/6 mice that previously received GFP^+^-OT-I cells, revealed that antigen-driven T cell expansion and IFN-γ production, was weaker in mice vaccinated with OVA-pulsed *Cish*^−/−^ GM-MØs (**Figure 1E-F**). The difference in T cell responses induced by the WT and *Cish*^−/−^ GM-MØs was not due to antigen uptake or processing, since both GM-MØs had similar uptake and processing capacity (**Figure 1G**). Moreover, when we bypassed antigen uptake/presentation by stimulating purified CD8^+^ T cells with anti-CD3/anti-CD28, *Cish*^−/−^ GM-MØs (at a ratio of MØs to T cells 1:1 and 1:2) were more potent at suppressing T cell proliferation and IFN-γ production than WT GM-MØs (**Figure 1H**). Thus, we conclude that CIS deficiency leads to the generation of MØs that strongly inhibit T-cell expansion.

### CIS deficiency leads to the generation of mouse and human MØs with reduced IL-12 production

We next investigated the mechanism underlying the lack of IFN-γ production by CD8^+^ T cells when co-cultured in presence of *Cish^−/−^* GM-MØs (**Figure 1F&H**). As IL-12 is a key cytokine promoting IFN-γ production (Henry et al., 2008), we measured IL-12 production by MØs upon TLR agonism. Compared to WT GM-MØs, IL-12 production by *Cish^−/−^* GM-MØs was substantially reduced when they were stimulated with CpG or LPS (**Figure 2A, Figure S4A**). WT and *Cish^−/−^* BM co-cultures highlighted that reduced IL-12 production by *Cish^−/−^* GM-MØs is cell-intrinsic (**Figure 2B**). In addition, we noted that IL-12 production was also reduced in *Cish^−/−^* GM-MØs stimulated with either PolyI:C or agonistic anti-CD40 antibody (**Figure S4B**). Interestingly, following CpG stimulation production of other cytokines (e.g. IL-6, IL-10 and TNF-α) were similar between WT GM-MØs and *Cish^−/−^* GM-MØs (**Figure S4C**), thus CIS deficiency in GM-MØs resulted in a selective impairment in IL-12 production. Further, the defect in IL-12 production by *Cish^−/−^* GM-MØs was also evident when they were derived from BM cultures across a range of GM-CSF concentrations (**Figure S4D**). In line with earlier reports (Helft et al., 2015; Sun et al., 2018), GM-DCs produced less IL-12 than GM-MØs and its production was CIS independent (**Figure S4E**). These results suggest that CIS plays a critical role in enabling IL-12 production by GM-MØs.

**Figure 2.**
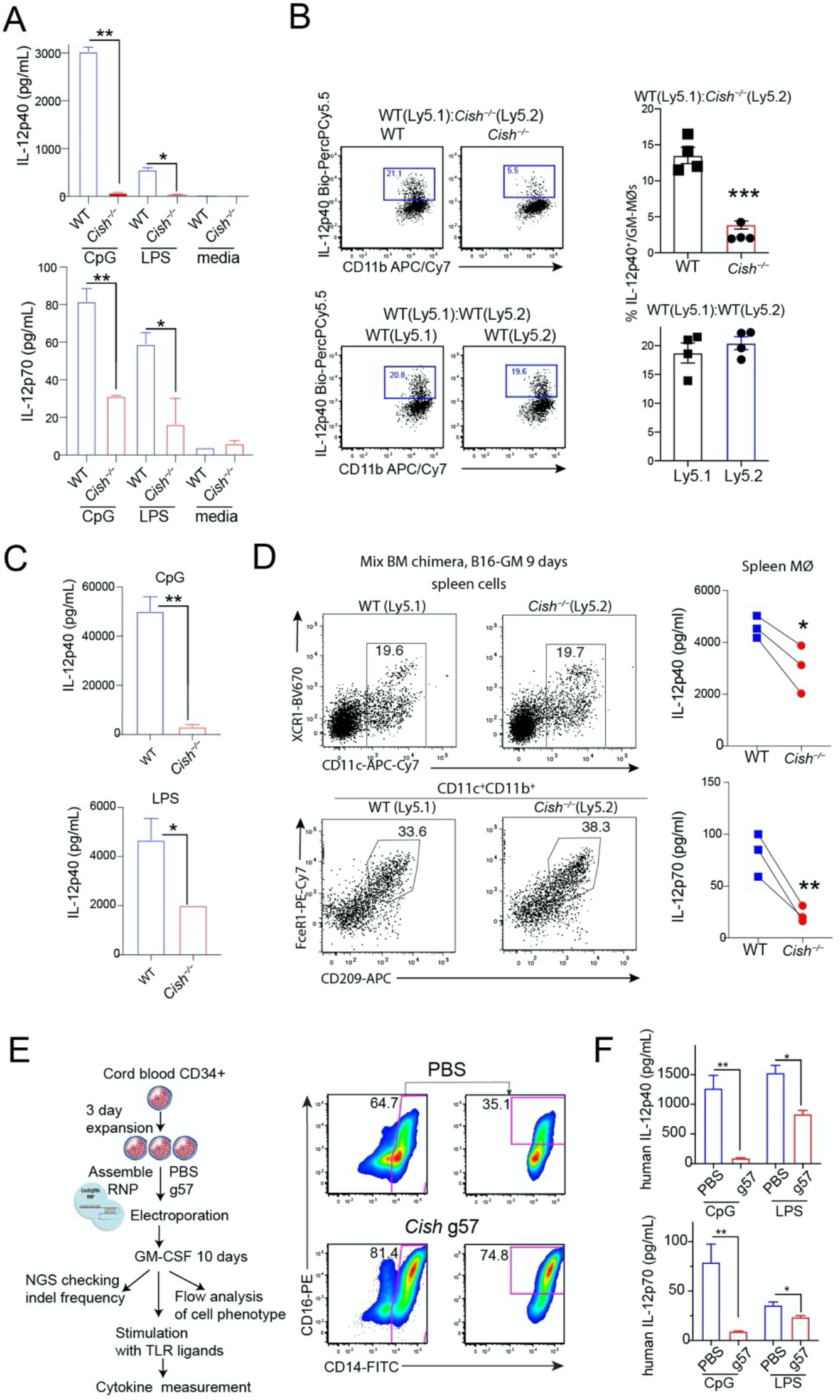
*Cish*^−/−^ MØs produce less IL-12. (**A**) IL-12 production by WT and *Cish*^−/−^ GM-MØs stimulated with GpG or LPS for 20 hrs. *P<0.05, **P<0.01 by multiple group comparison of ANOVA test. **(B**) Production of IL-12 by WT (Ly5.1) and *Cish*^−/−^ (Ly5.2) GM-MØs derived from co-cultures with BM cells from individual mice. Harvested cells were stimulated with CpG for 4 hrs. ***P<0.001 by t test. (**C**) IL-12 production by spleen cells from WT and *Cish*^−/−^ mice. Mice were engrafted with B16-GM for 9 days. Spleen cells were stimulated with CpG or LPS for 20 hrs. Bar graphs show mean ± SD of IL-12 concentration in culture supernatants. *P<0.05, **P<0.01 by t test. (**D**) IL-12 production by spleen MØs of WT and *Cish*^−/−^ mice. Mix bone marrow chimera mice reconstituted with both WT (Ly5.1) and *Cish*^−/−^ (Ly5.2) BM cells were engrafted with B16-GM for 9 days. Sorted MØs were stimulated with CpG for 20 hrs. Line graphs show IL-12 production by paired spleen MØs of WT and *Cish*^−/−^ mice from the same hosts. *P<0.05, **P<0.01 by t test. (**E&F**) IL-12 production by human GM-MØs with *Cish* deletion. Human cord blood CD34^+^ cells were transfected with Cas9 assemble RNP and *Cish* guide RNA. Transfected CD34^+^ cells were cultured with human GM-CSF (5 ng/ml) for 7 days. Cells were stimulated with CpG or LPS. FACS plots show resulted human GM-MØ population (**E**). Bar graphs show IL-12 production by human GM-MØs (**F**). *P<0.05, **P<0.01 by multiple group comparison of ANOVA test. Data are representative of three independent experiments.

The limited number of splenic MØs, (identified as CD11c^+^CD11b^+^CD209^+^FceR1^+^) probably due to low abundance of GM-CSF in unchallenged mice(Chow et al., 2016; Erlich et al., 2019), precluded us from corrobating the above findings in vivo. To circumvent this issue, we challenged wt or *Cish^−/−^* mice for 9 days with the B16 melanoma cell line producing GM-CSF(Dranoff et al., 1993). When splenocytes isolated from challenged WT and *Cish^−/−^* mice were stimulated with CpG or LPS, we found that IL-12 production was substantially reduced in *Cish^−/−^* splenocytes (**Figure 2C**). To evaluate the contribution of MØs to IL-12 production, we generated mixed bone marrow chimeras mice where lethally irradiated host (57BL/6-Ly5.1) reconstituted with both WT (Ly5.1) and *Cish^−/−^* (Ly5.2) BM cells. Following reconsititution, mice were then engrafted with B16-GM for 9 days. MØs of both WT and *Cish^−/−^* origin were isolated and stimulated with CpG for 20 hrs. Similar to in vitro generated GM-MØs, splenic *Cish^−/−^* MØs isolated from B16-GM bearing mice produced significantly less IL-12 than WT MØs (**Figure 2D**).

Finally, we investigated whether CIS impacts on IL-12 production by human MØs. To this end, human cord blood CD34^+^ cells were transfected by electroporation with Cas9 assembled in a ribonucleoprotein particle (RNP) with *Cish* guide RNA. The indel frequency for donor CD34^+^ cells with CIS RNP was 83% (based on next-generation sequencing). Transfected CD34^+^ cells were cultured with human GM-CSF (5 ng/mL) for 7-10 days. *Cish* guide and mock-transfected cells differentiated into CD14^+^CD16^+^ cells. Cell yields with *Cish* guide were slightly higher than mock-transfected. Notably, the percentage of CD14^+^CD16^+^ was higher in cultures with *Cish* guide (**Figure 2E**). When cells were stimulated with CpG or LPS, the production of IL-12 by cells targeted with *Cish* guide was substantially reduced when compared to the mock-transfected cells (**Figure 2E**). Taken together, these data point to a critical role for CIS in controlling the production of IL12 of both mouse and human MØs.

### CIS deficiency imprints MØs with M2 MØ characteristics

We showed above that *Cish^−/−^* MØs strongly inhibit T cell responses and produce less IL-12, compared to WT MØs. To gain insights into the molecular mechanisms accounting for these functional changes, we performed RNA sequencing (RNA-seq) on sorted GM-MØs and GM-DCs from 4 individual WT and *Cish*^−/−^ mice. Principal component analysis (PCA) revealed that CIS and cell lineage identity account for most differences between these samples. The impact of CIS deficiency was more prominent in GM-MØs than in GM-DCs as evidenced by the increased separation of the MØ groups in the PCA plot (**Figure 3A**), and the higher number of differentially expressed genes between the *Cish*^−/−^ and WT (1,029 and 713 DEGs for GM-MØs and GM-DCs, respectively) (**Supplemental Table 1**). As MØs are the dominant cell type generated in GM-CSF stimulated BM culture and more profoundly impacted by CIS deficiency, we chose MØs for further detailed analysis. Gene ontology analysis highlights many differences between WT and *Cish*^−/−^ GM-MØs that might correlate to their differences in functions, cell cycle and metabolism (**Figure S5A**).

**Figure 3.**
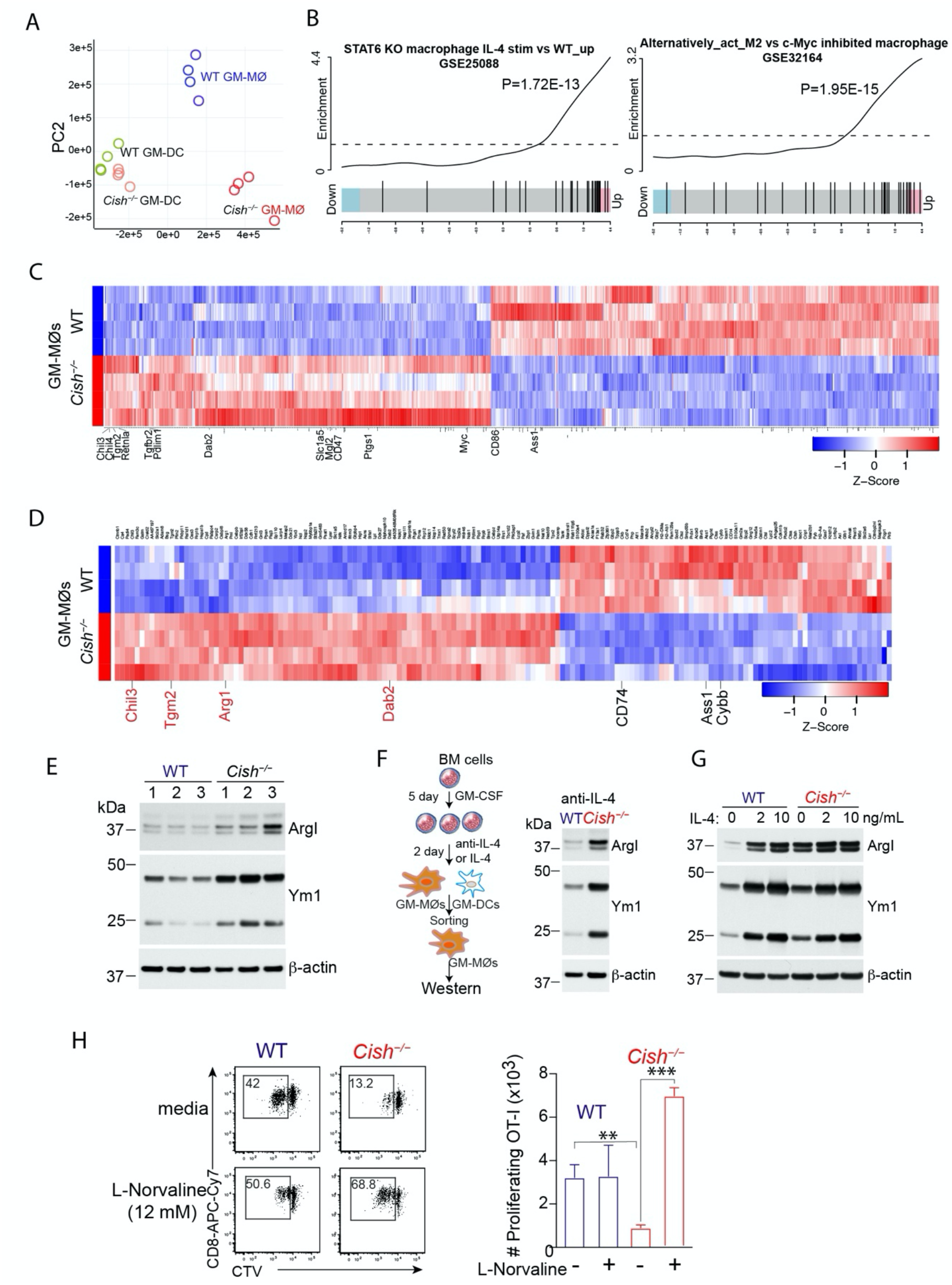
*Cish*^−/−^ MØs display characteristics of M2 MØs. (**A**) Principal component analysis (PCA) plot of TPM values over dimensions 1 and 2 with samples from 4 individual mice per group colored by groups of GM-MØs and GM-DCs. (**B**) Gene set enrichment analysis barcode plot comparing the expression of upregulated genes in *Cish^−/−^* GM-MØs, with two previously reported M2 MØs signature gene sets. (**C**) *Cish*^−/−^ GM-MØs expressing higher levels of many M2 markers. Heatmap shows DEGs with M2 signature genes indicated in red. (**D**) Proteomic analysis of GM-MØs derived from 4 individual WT and *Cish*^−/−^ mice. Heatmap show expression of differentially regulated proteins by two types of GM-MØs, with M2 proteins indicated in red (LFC>0.6, FDR<0.05). (**E**) Western immunoblot analysis of Arg1 and Ym1 by WT or *Cish*^−/−^ GM-MØs derived from 3 individual donors. (**F**& **G**) BM cells were cultured with GM-CSF for 5 days, anti-IL-4 Ab or recombinant IL-4 were then added. At 2 more days, GM-MØs were sorted and analysed for Arg1 and Ym1 expression by Western immunoblot. (F) Cultures with or without anti IL-4 Abs; (G) Cultures with or without recombinant IL-4. (**H**) CTV-labelled OT-1 CD8^+^ T cells were co-cultured with GM-MØs and stimulated OVA in the absence or presence of L-Norvaline. Dot plots show OVA-stimulated proliferation of CTV-labelled CD8^+^ T cells. Bar graphs show numbers of proliferating CD8^+^ T cells. *P<0.05, **P<0.01, **P<0.001 by multiple group comparison of ANOVA test.

GM-CSF is thought to bias MØs polarization towards the pro-inflammatory M1-like state (Fleetwood et al., 2007; Martinez and Gordon, 2014), and thus we expected that CIS deficiency would strengthen GM-CSF signalling and further increase M1 polarization. Instead, analysis of RNAseq data indicates that the genes upregulated in *Cish^−/−^* GM-MØs were positively correlated to gene signatures derived from IL-4 induced M2 MØs (GSE25088) (Szanto et al., 2010) and GSE32164 (Pello et al., 2012) (**Figure 3B**). Next, we analyzed genes known to be associated with MØ functional polarization among DEGs in *Cish^−/−^* GM-MØs (Martinez et al., 2006; Martinez et al., 2013) (**Supplemental Table 2**). Over 64% (35/56) of the M2-associated DEGs (**Supplemental Table 2**) were up-regulated in *Cish^−/−^* GM-MØs (**Figure 3C**), including prototypic genes such as *Chil3 (Ym1), Chil4, Retnla (Fizz1) and Tgm2* while over 85% (26/30) (**Supplemental Table 2**) of the M1 associated DEGs were down-regulated in *Cish^−/−^* GM-MØs, thus suggesting that CIS deficiency leads GM-CSF induced MØ toward a M2-like phenotype.

To support our transcriptomic analysis, we also performed label-free quantitative proteomics analysis from GM-MØs derived from 4 wt and *Cish^−/−^* mice (**Supplemental Table 3**). In accordance with the transcriptional data, we found that several typical M2 associated proteins were up-regulated in *Cish^−/−^* GM-MØs including Arg1, Chil3, Tgm2 and Dab2 (**Figure 3D**). In contrast, proteins that were down-regulated in *Cish^−/−^* GM-MØs included M1 associated proteins such as CD74, Ass1 and Cybb (**Figure 3D**). Overall, both RNAseq and proteomic analyses revealed that the loss of CIS results in GM-MØs developing M2-like features.

Next, we selected two commonly used M2 markers Arg-1 and Ym1 for further validation by Western blot. *Cish^−/−^* GM-MØs sorted from BM cultures of 3 individual mice all had higher expression of Arg-1 and Ym1 than WT GM-MØs (**Figure 3E**), thus corroborating the results hereabove. In an attempt to convert *Cish^−/−^* GM-MØs into M1 like MØs, we stimulated WT and *Cish^−/−^* GM-MØs with IFN-γ and LPS, potent M1 inducing cytokines. While IFN-γ and LPS stimulation of GM-MØs reduced expression of Arg-1 and Ym1 in both WT and *Cish^−/−^* GM-MØs, the latter retain a substantial higher amount of these M2 markers (**Figure S5B**).

IL-4 is a prototypic cytokine that differentiates M2 MØs (Martinez et al., 2013). To investigate whether M2 characteristics of *Cish^−/−^* GM-MØs are influenced by endogenous IL-4, neutralizing Ab to IL-4 was added to BM cell culture with GM-CSF at day 5. Even when GM-MØs were isolated from co-culture of WT and *Cish^−/−^* BM cells, *Cish^−/−^* GM-MØs still had higher expression of YM-1 and Arg1 (**Figure 3F**), suggesting that induction of M2 phenotype in *Cish^−/−^* GM-MØs is unlikely to be directly due to IL-4. Further, addition of exogenous IL-4 in the late stage of cell differentiation increased the expression of Arg1 and Ym1 in GM-MØs but this increase was comparable between WT and *Cish^−/−^* genotypes (**Figure 3G**).

Finally, we examined the functional consequence of the high level of Arg1 in *Cish^−/−^* GM-MØs. Arginine availability is key to an optimal T cell immune response (Rodriguez et al., 2007). As the Arginase 1 inhibitor L-Norvaline enhances T cell proliferation in the presence of antigen-presenting cells(Bronte et al., 2003), we investigated whether inhibition of Arg-1 activity in *Cish^−/−^* GM-MØs could relieve the suppressive activity of *Cish^−/−^* GM-MØs (**Figure 1D-H**). When L-Norvaline was added during T-cell stimulation, the proliferation of OT-1 was greatly enhanced in cultures with *Cish^−/−^* GM-MØs, but not WT GM-MØs (**Figure 3H**). Thus, we contend that M2 molecular imprinting in the absence of CIS indeed leads to a gain of suppressive function which can be partially relieved by inhibiting Arginase 1.

### Enhanced STAT5 activation in the absence of CIS contributes to the development of M2-like MØs

Next we interrogated the signaling events leading to the development of M2-like features in the absence of CIS. Isolated GM-MØs restimulated with GM-CSF induced expression of CIS mRNA and protein and STAT5 activation in GM-MØs (**Figure 4A**&**B**). Further, CIS deficiency in GM-MØs led to enhanced and prolonged STAT5 activation after GM-CSF restimulation (**Figure 4C**), but no differences in PI3K and MAPK pathways was detected **(Figure 4D).**

**Figure 4.**
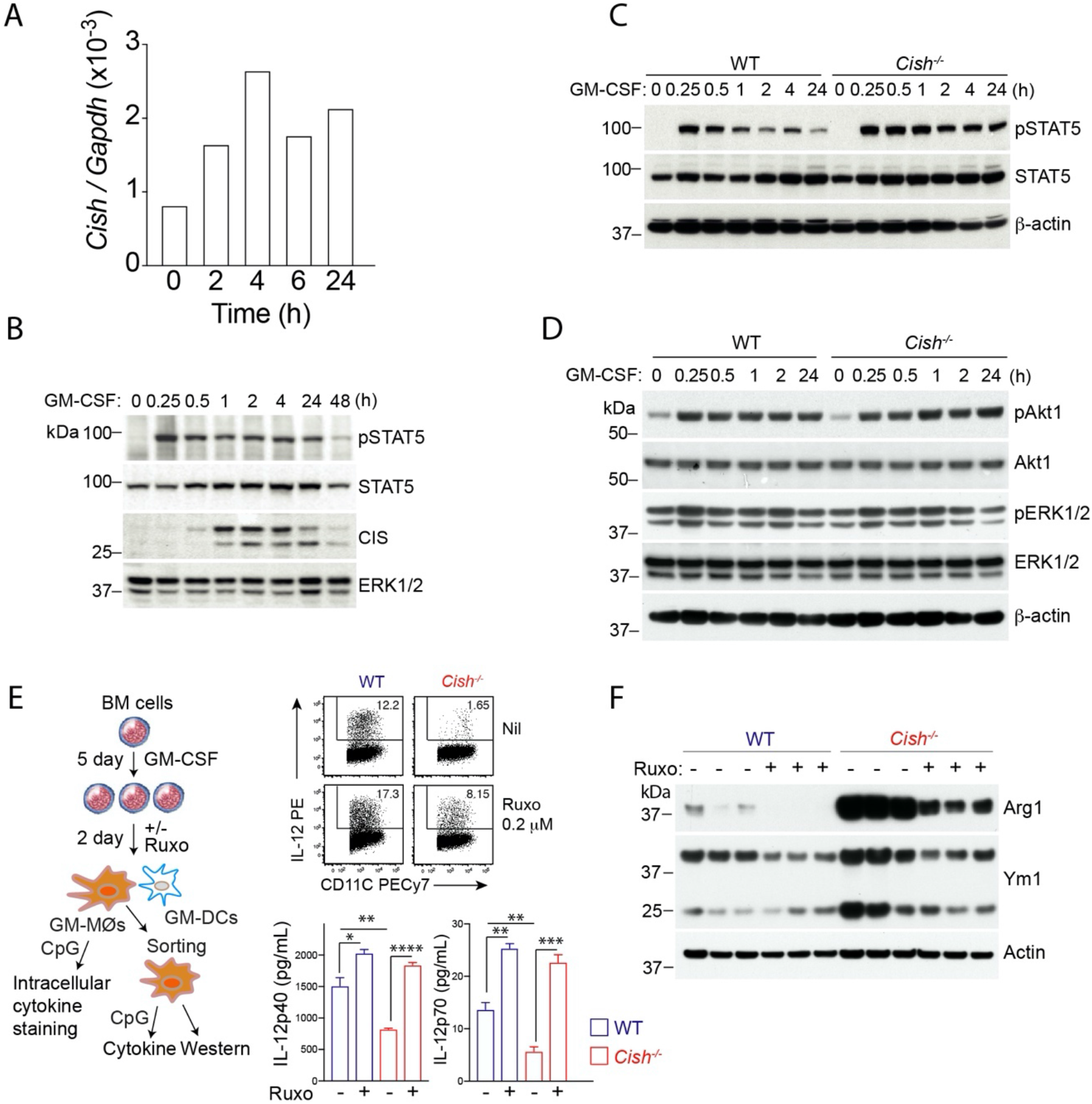
*Cish*^−/−^ GM-MØs have enhanced STAT5 activation, contributing to a M2-like phenotype. **(A&B)** Purified GM-MØs were stimulated in vitro with recombinant GM-CSF (50 ng/ml) for indicated times in RPMI1640, supplemented with antibiotics and 10% heat-inactivated FCS. Cells were washed in cold PBS and pelleted. Induction of *Cish* mRNA relative to house keeping gene GAPDH in GM-MØs upon GM-CSF stimulation was determined by RT-qPCR from extracted RNA samples (**A**). Induction of CIS protein and STAT5 activation in GM-MØs upon GM-CSF restimulation were determined by Western immunoblot (**B**). (**C&D**) Purified WT and *Cish*^−/−^ GM-MØs were stimulated in vitro with recombinant GM-CSF (50 ng/ml) for indicated times. Cells were lysed for Western immunoblot against STAT5 activation (**C**); Akt2 and ERK activation (**D**). (**E&F**) BM cells were cultured with GM-CSF for 5 days and were supplemented with or without JAK inhibitor ruxolitnib (RUXO) (0.2 μM) for 2 days. IL-12 production by GM-MØs upon CpG stimulation was evaluated by intracellular cytokine staining and secretion **(E)**. *P<0.05, **P<0.01, ***P<0.001, ****P<0.0001 by multiple group comparison of ANOVA analysis. Data are representative of two independent experiments. (**F**) Purified WT and *Cish*^−/−^ GM-MØs were also analysed for Ym1 and Arg1 expression by Western Blot. Data derive from 3 individual mice.

To investigate whether enhanced JAK/STAT5 signaling is responsible for some of the functional alterations observed in *Cish^−/−^* GM-MØs, we tuned down GM-CSF signaling using the JAK inhibitor ruxolitinib (RUXO, 0.2 μM). Addition of RUXO to the cultures in the last two days of 7-day culture was able to rescue the capacity of *Cish^−/−^* GM-MØs to produce IL-12 production following CpG stimulation, and IL-12 production by wt GM-MØs was further enhanced by RUXO (**Figure 4E**). This suggests that decreasing JAK/STAT5 signalling output in GM-MØs, normaly mediated by CIS, is critical for optimal IL-12 production. In addition, we examined the expression of the M2 markers Arg1 and Ym1 following the inhibition of the JAK/STAT5 pathway. As previously shown **(Figure 2)**, GM-MØs derived from *Cish^−/−^* BM had a higher expression of Arg and Ym1, which was greatly reduced following RUXO addition to the cultures (**Figure 4F**). Taken together, these data suggest that the lack of CIS in GM-MØs results in an increased JAK/STAT5 signalling pathway leading to the development of M2-like features and reduced IL-12 production.

### Downregulation of IRF8 by enhanced STAT5 activation contributes to the development of M2-like MØs

To define the molecular consequences of the sustained JAK/STAT5 activation observed in *Cish^−/−^* GM-MØs, we interrogated our transcriptomics dataset. We focused our analysis on transcription factors known to regulate M1/M2 cell fate, and in particular members of the interferon responsive factor (IRF) that are known to be critical for MØ polarization (Gunthner and Anders, 2013). Quantitation of the mRNA for the 9 members of this family revealed that *Irf8* mRNA transcripts were substantially reduced *Cish^−/−^* GM-MØs compared to WT GM-MØs, as opposed to the other family members whose transcripts were not altered by CIS deficiency (**Figure 5A**). To substantiate these findings we crossed *Cish^−/−^* mice with *Irf8^Gfp^* (Wang et al., 2014), allowing us to measure IRF8 expression at a single-cell level. GM-CSF reduced expression of IRF8 in GM-MØs in a dose dependent manner (**Figure 5B**). Comparison of GM-MØs derived from *Irf8^Gfp^/WT* and *Irf8^Gfp^*/*Cish^−/−^* mice confimed the reduced expression of IRF8 in GM-MØs lacking CIS (**Figure 5B**). Consistent with the above, the expression of IRF8 was substantially reduced in MØs isolated from *Irf8^Gfp^*/*Cish^−/−^* mice that had been challenged with B16-GM tumors 9 day previously, compared to similarly treated *Irf8^Gfp^*/*WT* mice (**Figure 5C**). We next investigated whether attenuation of JAK/STAT signaling could rescue *Irf8* transcription in *Cish^−/−^* GM-MØs. To this end, *Irf8^Gfp^/Cish^−/−^* BM progenitors were cultured in presence of GM-CSF for *Irf8^Gfp^/Cish^−/−^* and on day 5 RUXO was added or not to the culture. On day 7, we noted that *Irf8^Gfp^/Cish^−/−^* GM-MØs treated with RUXO expressed higher IRF8-GFP than untreated *Cish^−/−^* GM-MØs (**Figure 5D**). Similar observations were made for GM-DCs (**Figure 5D**). This suggests that the sustained JAK/STAT5 activation observed in *Cish^−/−^* GM-MØs impairs is detrimental for IRF8 expression.

**Figure 5.**
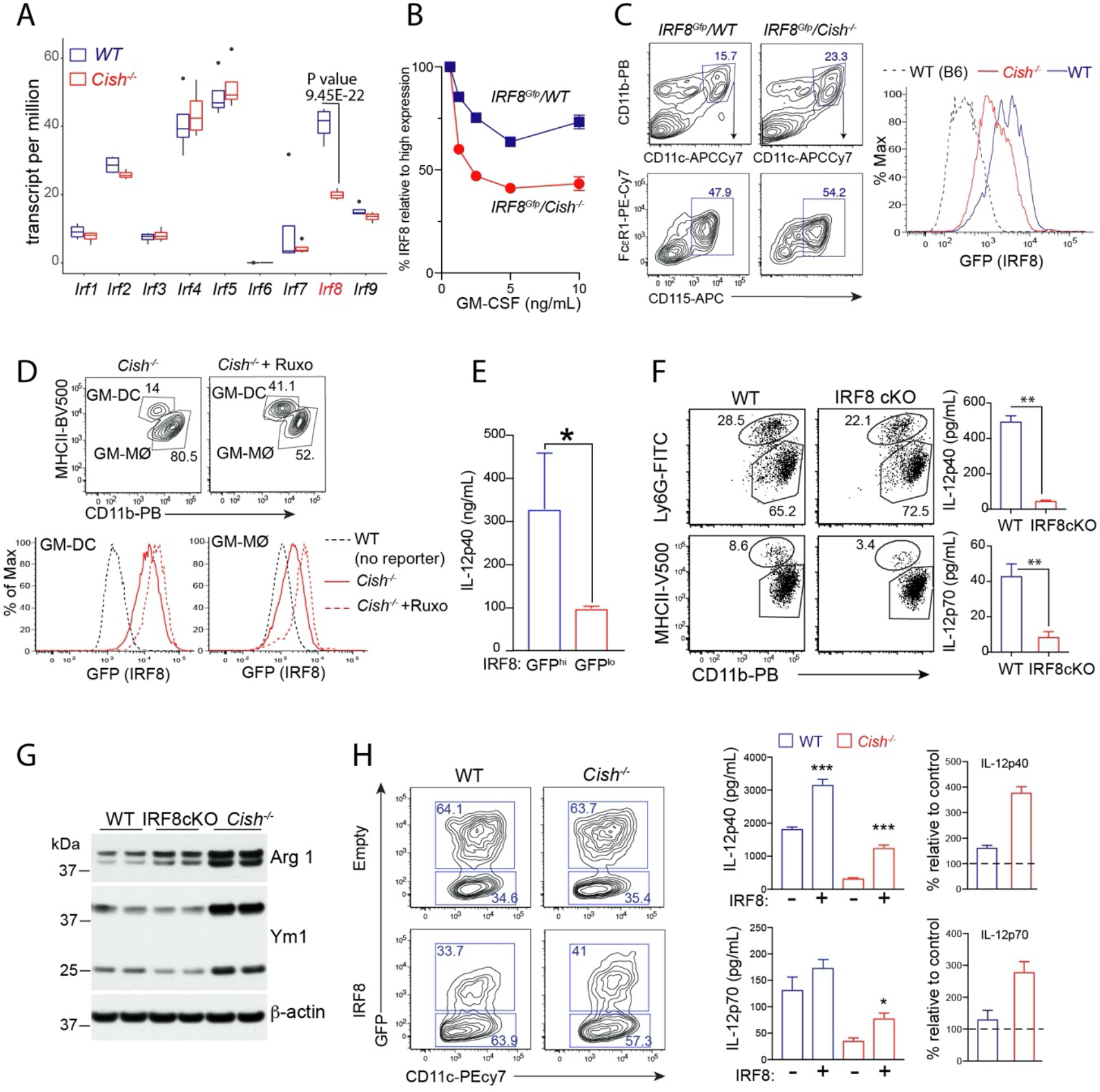
*Cish*^−/−^ MØs have lower expression of IRF8, resulting in a M2-like phenotype. (**A**) Expression of the Interferon regulatory factors (IRFs) by WT and *Cish*^−/−^ GM-MØs was analysed by RNAseq. Boxplot shows IRF expression of WT and *Cish*^−/−^ GM-MØs. (**B**) BM cells from *IRF8^Gfp^* mice with or without *Cish* deficiency were cultured with different concentrations of GM-CSF. Heatmap shows MFI of *IRF8^Gfp^* by GM-MØs. (**C**) *IRF8^Gfp^* mice with or without *Cish* deficiency were engrafted with B16-GM cells. Isolated spleen MØs were evaluated for expression of IRF8 reporter. Negative controls (dot line) are MØs from B6 mice. (**D**) *IRF8^Gfp^*/*Cish*^−/−^ BM cells were cultured in the presence of GM-CSF for 5 days. Cells were then cultured with Jak Inhibitor RUXO for further 2 days and GFP expression by GM-MØs and GM-DCs was evaluated. (**E**) GM-MØs from *IRF8^Gfp^* mice sorted into top 50% (GFP^hi^) and bottom 50% (GFP^lo^) based GFP (IRF8) expression. Sorted GM-MØs were then stimulated with CpG for 20 hours and IL-12 concentration from cultured supernatants measured by ELISA. *P<0.05 by t test. (**F&G**) GM-MØs from *CD11c-cre-IRF8^fl/fl^* (IRF8cKO) and WT mice were stimulated with CpG for 20 hrs. IL-12 production was then measured and presented (F). **P<0.01 by t test. Sorted GM-MØs cultured fro BM cells of IRF8cKO, WT and *Cish^−/−^* mice were also analysed for expression of Arg1 and Ym1 by Western immunoblot analysis (G). (**H**) BM progenitor cells from WT and *Cish^−/−^* were transduced with either Empty/GFP, or IRF8/GFP encoding vector and cultured with 10 ng/ml GM-CSF for 5 days. Gated represents the frequency of transduced GM-MØs (GFP^+^). Sorted transduced GM-MØs were stimulated with CpG for 20 hrs. IL-12 in culture supernatants were presented as bar graphs. Right panels show % increase in IL-12 production by cells with IRF8/GFP encoding vector relative to cells with empty/GFP encoding vector of the same genotype. Data are representative of 3 independent experiments. *P<0.05, ***P<0.001, by multiple group comparison of ANOVA analysis.

We then explored whether the downregulation of IRF8 contributed to the impaired IL-12 production observed *Cish^−/−^* GM-MØs. The first line of support came from the experiment with the *Irf8^Gfp^* reporter mouse. When GM-MØs from *Irf8^Gfp^* mice were arbitrarily segregated into two populations based on GFP intensity, we found that GFP^hi^ GM-MØs produced significantly higher levels of IL-12 than GFP^lo^ GM-MØs (**Figure 5E**). This suggested that in GM-MØs subtle changes in IRF8 expression could result in functional differences. We then tested whether the lack of IRF8 impacted the capacity of GM-MØs to produce IL-12. To circumvent the issue of the defective myelopoeisis observed in germline IRF8 KO (Kurotaki et al., 2014), we used *CD11c-Cre*/*Irf8*^fl/fl^ (IRF8cKO) mice to generate GM-MØs. In such a system, the compartment of Ly6G^+^ granulocytes was similar between WT and IRF8cKO while GM-DCs were slightly reduced in IRF8cKO (**Figure 5F**). Consistent with our earlier observations, GM-MØs derived from IRF8cKO mice had reduced IL-12 production following CpG stimulation when compared to their WT counterparts (**Figure 5F**). As *Cish^−/−^* GM-MØs expressed higher levels of number of M2 markers including Arg1 and Ym1, we evaluated whether IRF8cKO GM-MØs would show similar features. The expression of Arg1 by IRF8cKO was consistently higher in IRF8 deficient GM-MØs when compared to WT GM-MØs. However, the phenotype of IRF8cKO GM-MØs was somewhat less pronounced than that of *Cish^−/−^* GM-MØs (**Figure 5G**). In an attempt to rescue IL-12 production by *Cish^−/−^* GM-MØs, we over-expressed IRF8 in GM-MØs using a retroviral system. IRF8 overexpression in *Cish^−/−^* GM-MØs, substantially increased IL-12 production following CpG stimulation, suggesting that the reduced Irf8 expression observed in *Cish^−/−^* GM-MØs impaired their capacity to produce IL-12 (**Figure 5H**). Collectively, our data suggest that CIS deficiency leads to increased STAT5 activation, resulting in the downregulation of IRF8 and therefore hindering their polarization into M1 like cells.

### IRF8 regulates the transcriptional programing of MØ polarization

Our results here above points to a critical role for CIS in maintaining an adequate IRF8 concentration which ultimately controls MØ polarization. To gain insight into the potential instructive role of IRF8 regulation in controlling GM-MØ polarization, we performed Cut&Tag sequencing (Kaya-Okur et al., 2019) to identify the genes directly targeted by IRF8 in GM-MØ. To facilitate our pulldown strategy, we derived GM-MØs from our *Irf8^Gfp^* WT and *Irf8^Gfp^*/*Cish^−/−^* mice in which IRF8 is a fusion protein with GFP, and the pull-down was performed with an anti-GFP Ab (**Figure S6A&B**). Using this strategy, we identified 12,142 binding sites in the genome of WT GM-MØs occupied by IRF8, and 10,270 binding sites in the *Irf8^Gfp^*/*Cish^−/−^* GM-MØs (**Figure 6A**). Comparison of our genome-wide IRF8 binding data set with our RNAseq allowed us to identify putative IRF8 regulated genes. For genes positively regulated by CIS (down in the *Cish^−/−^* GM-MØs), 34% of them displayed at least one IRF8 binding site in proximity of their transcritpion starting site (TSS) while for genes negatively regulated by CIS (therefore up in the *Cish^−/−^* GM-MØs), 28% of them displayed at least one IRF8 binding site in proximity of their TSS (**Figure 6B**). As we reasoned that CIS deficiency in GM-MØs leads to a perturbed polarization potential, in part due to the lack of IRF8 upregulation, we focused on the genes associated with MØ polarization (**Supplemental Table 1**). For M1-associated genes, 40% of the M1 genes positively regulated by CIS (*Ace, Cd74* and *Cyyb*) were bound by IRF8, while only 3% of the M1 genes negatively regulated M1 by CIS (*Dusp6*) were bound by IRF8 (**Figure 6C**). On the other hand, of the 56 M2-signature genes bound by IRF8, 30% were negatively regulated by CIS (*Ahr, Chil4* and *Myc*) and 14% were positively regulated by CIS (*F13a1* and *Msr1*) (**Figure 6D**). Taken together these observations support that CIS modulation of IRF8 plays a key role in the regulation of M1 like state.

**Figure 6.**
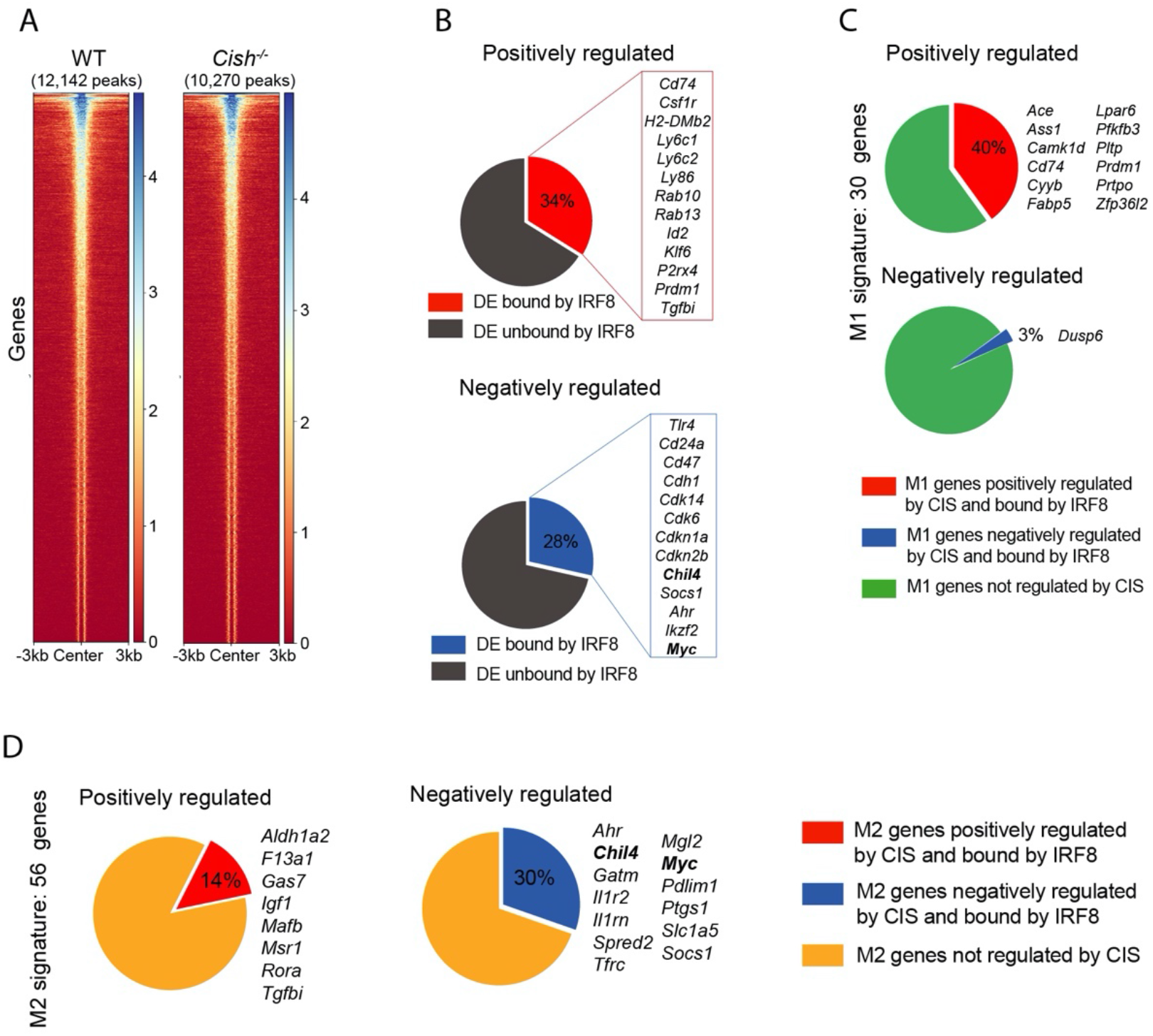
IRF8 contains binding sites for genes associated MØ polarization. (**A**) Heatmap showing the IRF8 binding in WT and *Cish^−/−^* GM-MØs, number below shows the binding site for IRF8 in each genotype GM-MØs. (**B**) DEGs of RNA-seq displaying at least one IRF8 binding site in their locus. The peaks were assigned to the closest gene. Top: genes positively regulated by CIS (therefore down in the *Cish^−/−^* GM-MØs); Bottom: genes negatively regulated by CIS (therefore up in the *Cish^−/−^* GM-MØs). (**C**) M1 associated DEGs displaying IRF8 binding site in their locus. Top: M1 associated genes positively regulated by CIS; Bottom: M1 associated genes negatively regulated by CIS. (**D**) M2 associated DEGs displaying IRF8 binding site in their locus. Top: M2 associated genes positively regulated by CIS; Bottom: M2 associated genes negatively regulated by CIS.

### *Cish*^−/−^ MØs increase Th2 response and the severity of allergic asthma

To examine if the M2-like MØ phenotype observed in the absence of CIS evoked a Th2 response in vivo, we transferred OVA-pulsed WT and *Cish^−/−^* GM-MØs into B6 mice and measured the production of IL-4 and IFN-γ from restimulated splenocytes after 6 days. We noted that level of IL-4 produced was substantially increased in mice that received OVA-pulsed *Cish^−/−^* MØs compared to mice immunized with OVA-pulsed WT MØs, while IFN-γ levels were reduced (**Figure 7A**). Ex vivo evaluation of OVA-stimulated CD4^+^ T cells confirmed that *Cish^−/−^* MØs induced more IL-4-producing cells (**Figure 7B**). Next, we compared the impact of transferred OVA-pulsed MØs of WT and *Cish^−/−^* origin on the induction of a Th2 allergic inflammation (Chavez-Galan et al., 2015). To this end, we immunized WT mice with OVA-pulsed WT MØs or *Cish^−/−^* MØs and then challenged with nebulized OVA 4 weeks later (**Figure 7C**). Upon challenge, lungs from mice that were pre-immunised with OVA-pulsed *Cish^−/−^* MØs contained a substiantial increased in inflammatory immune cells compared to mice that received OVA-pulsed WT MØs (**Figure 7D**). In line with increased inflammation, mice pre-immunised with OVA-pulsed *Cish^−/−^* MØs showed a high prevalence of PAS^+^ mucous-producing cells symptomatic of the development of an allergic airway disease. Additionaly, the characterisation of the cytokine content of bronchoalveolar lavage fluid (BALF), revealed that inflammatory cytokines such as IL-4, IL-6 and IL-5 were significnatly higher in the BALF of mice than mice preminnunised pre-immunised with OVA-pulsed *Cish^−/−^* compared to mice premimmunised with OVA-pulsed WT MØs (**Figure 7E-G**). Notably, when IL-4 to IFN-γ ratio from the same sample was calculated to reflect a potential Th2 bias, BALF from the mice immunized with OVA-pulsed *Cish*^−/−^ MØs contained preferentially increased Th2 cytokine IL-4 (**Figure 7H**). Taken together, these data suggest that CIS deficiency results in the polarization of MØs featuring a strong capacity in inducing a robust Th2 response, that exacerbates the allergic asthma response. Thus, CIS controls Th differentiation through the control of MØ polarization independent of the reported role of CIS in controlling intrinsically T cell polarisation(Yang et al., 2013).

**Figure 7.**
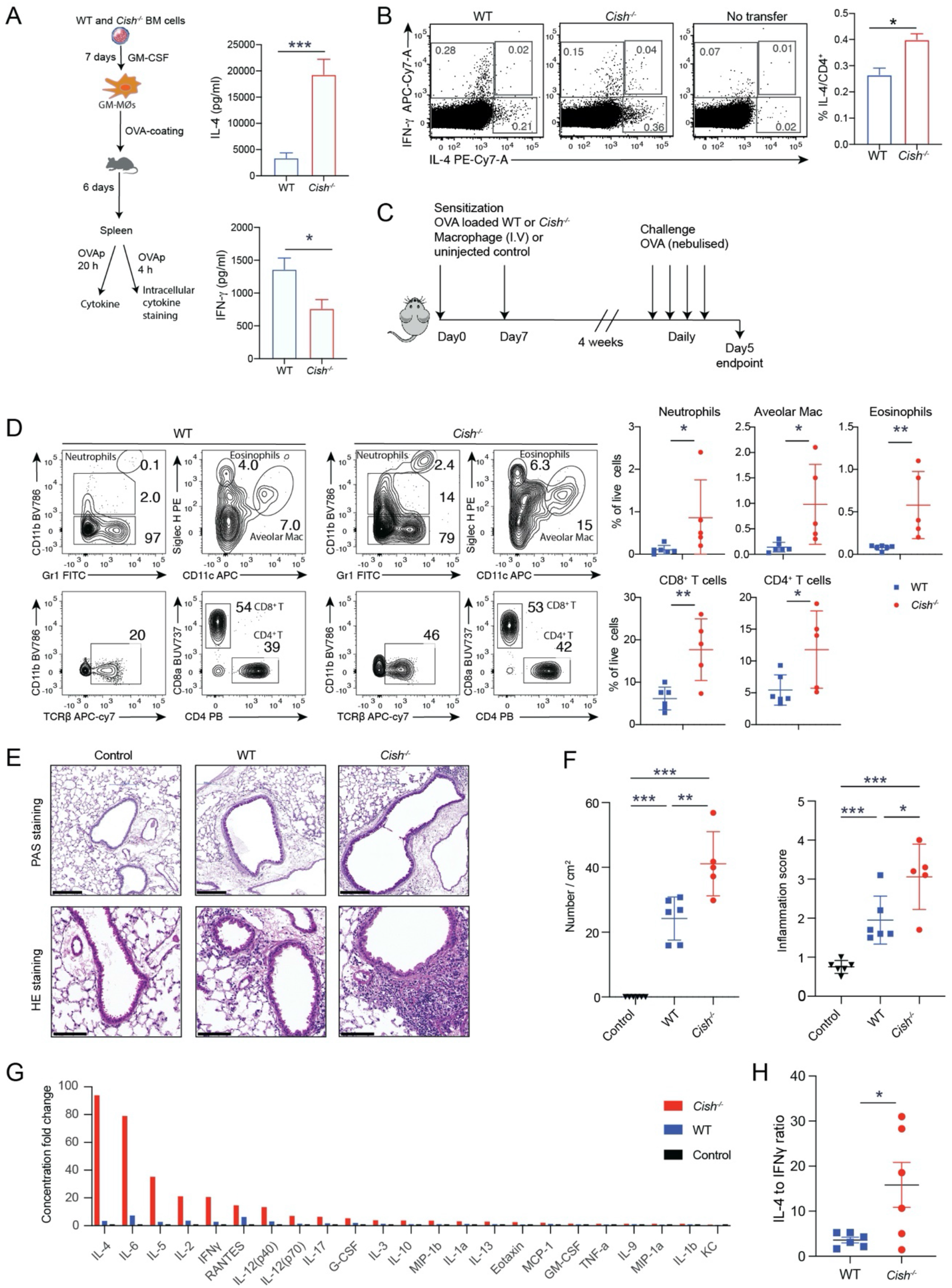
*Cish^−/−^* MØs promote Th2 responses and exacerbates allergic asthma. **(A&B)** Schematic plot shows the experimental approach for assessing the immune response induced by WT or *Cish*^−/−^ GM-MØs. Six days after transfer of OVA-coated GM-MØs, harvested splenic cells from 3 individual mice were cultured with OVA for 20h. The supernatant was harvested for cytokine assay. Bar graph shows the concentration of IL-4, IFN-γ and IL-17A (**A**). (**B**) Splenic cells were pulsed with OVA_323-339_ for 4 hours. The proportion of of IL-4 and IFN-γ-producing CD4^+^ Tcells were tested via intracellular staining. Dot plots show the IL-4 and IFN-γ expression in CD4^+^ T cells. Bar graph shows % IL-4^+^ CD4^+^ T cells. *P<0.05 by t-test. **(C)** Experimental protocol of OVA-induced inflammation. **(D)** Contour plots show the gating strategy for neutrophils, alveolar macrophage, eosinophils, and T cells in bronchoalveolar lavage; Scatter plots show mean +/−SEM of different immune cell populations from individual mice treated as in (**C**). **(E)** Representative PAS and HE staining of the airway of the mice indicated in (**C**). Scale bar= 200μm. **(F)** Quantification of PAS^+^ cells and inflammation score. Bar graphs show mean+/−SEM of indicated populations. Each dot represents one mouse. **(G)** Cytokine in bronchoalveolar lavage fluid of the mice treated as in (**C**), were analysed by Bio-plex assay. Bar graph shows fold change over mean level in control mice. (**H**) Ratio of IL-4 to IFN-γ level from the same samples. Each dot represents one mouse **P<0.01, ***P<0.001 by t-test.

## Discussion

A central finding from our study is that CIS has a marked impact on GM-CSF-induced MØ polarization. In its absence, and in microenvironements where GMCSF was abundant, MØs became immunosuppressive, produced less IL-12, and promoted a Th2 immune response. In contrast to IL-4 induced M2 polarization that depends on the signaling axis of STAT6 and IRF4(Satoh et al., 2010). Our data demonstrate that CIS deficiency induced M2-like polarization following GM-CSF exposure, through sustained activation of STAT5 and downregulation of IRF8. We further showed that IRF8 regulates the gene expression program governing MØ polarization. Thus, we propose that CIS may represent a key intrinsic regulatory mechanism responsible for the functional polarization of MØs.

At the cellular level, both human and mouse GM-CSF treatment in vitro has been shown to promoted the differentiation of proinflammatory M1-like MØs (Fleetwood et al., 2007; Verreck et al., 2004). Similarly, in vivo, GM-CSF has been shown to promote the pathogenicity of MØs in murine models of autoimmunity associated with production of proinflammatory cytokines(Croxford et al., 2015; Ko et al., 2014). Nervetheless, GM-CSF has also been associated with the development of suppressive M2-like MØs in various settings(Bernasconi et al., 2010; Bronte et al., 1999; Huen et al., 2015). How GM-CSF drives these contrasting cellular fates of MØs is still largely unknown. A simple interpretation for the dichotomy of MØ polarization under GM-CSF stimulation is a model of ligand abundance mediated polarisation, as the development of suppressive M2-like MØs is largely associated with high GM-CSF availability (Bronte et al., 1999; Huen et al., 2015). At a molecular level, heightened STAT5 activation under GM-CSF stimulation had been linked to development of M2-like MØs (Huen et al., 2015; Zhang et al., 2020). Accordingly, neutralization of GM-CSF or STAT5 inhibition can reverse M2 polarization. Our study identifies another layer of control independent of GM-CSF abundance. Here, we report that CIS acts as a cell-intrinsic rheostat in controlling STAT5 activation independently of the activation of PI3K or MAPK. CIS deficiency in GM-MØs led to heightened and sustained STAT5 activation that favored M2-like MØs differentiation. Fittingly, attenuation of upstream signalling with a JAK inhibitor increased IL-12 production and reduced expression of Arg-1 and YM-1 by *Cish^−/−^* MØs.

Here we revealed a CIS-censored signalling axis of GM-CSF/STAT5/IRF8 in regulating MØ polarization. In our study, GM-CSF signalling in *Cish^−/−^* MØs leads to lower expression of IRF8 while JAK inhibition in *Cish^−/−^* MØs reduced STAT5 and increased IRF8 expression. STAT5 activation has been shown previously to suppress induction of *Irf8* transcription in plasmacytoid DCs (Esashi et al., 2008). As IRF8 has been reported as a critical positive regulator of IL-12 production (Holtschke et al., 1996; Masumi et al., 2002), a link between its downregulation and the reduced production of IL-12 observed in *Cish^−/−^* MØs seemed to offer a rationale for the functional impairment observed in these cells. Earlier studies provided circumstantial evidence to support the idea that the downregulation/silencing of IRF8 could represent a defining moment in favoring the development of M2-like MØs through the increased expression of M2 MØs genes (Ototake et al., 2021; Twum et al., 2019; Waight et al., 2013). Our data demonstrated that IRF8 bound in the vicinity of many genes that have been associated with MØ polarization, suggesting a critical instructive role in controlling M1 like features. IRF8 is unlikely to be the sole downstream target of GM-CSF/STAT5 pathway leading to M2 polarization.

Although M2-like MØs are known for their ability to promote Th2 differentiation(Mills et al., 2000), the molecular mechanisms responsible for this process are less understood. It has been previously reported that the M2 signatire genes Ym1/2, can directly influence Th2 development (Arora et al., 2006; Cai et al., 2009), with an anti-Ym1 antibody significantly reducing both GATA-3 expression and Th2 cytokine production by CD4^+^ T cells (Arora et al., 2006; Cai et al., 2009). Relevant to in vivo Th2 responses, *Cish^−/−^* GM-MØs expressed significantly higher levels of chemokines associated with Th2 immunity (Ozga et al., 2021), especially IL-4, compared to WT GM-MØs. Our transcriptomic and proteomic approached highlighted the high expression of TGM2, Arg1 and FIZZ1 in *Cish^−/−^* GM-MØs (**Figure 3**), all of which have been shown to play a role in allergic asthma (Abdelaziz et al., 2020). Thus, we contend that CIS expression in myeloid cells has a prominent role in regulation of Th2 immunity, independently of the reported role for CIS in T cells in promoting Th2 differentiation (Yang et al., 2013).

Together, we revealed here that CIS acts as a brake on GM-CSF signaling, and is critical for MØ polarization. By dissecting the molecular and functional nature of *Cish^−/−^* MØs, we provide significant insight into how GM-CSF shapes MØ polarization. We argue that CIS expression in myeloid cells may be beneficial in allergic response and anti-tumor immunity by curtailing GM-CSF signaling to prevent M2 polarization and pathogenic Th2 responses.

## Materials and Methods

### Mice

C57BL/6 (B6, WT), Ly5.1, Ly5.1/Ly5.2 F1, *Rag1^−/−^/Il2rγ^−/−^, Cish^−/−^*(Palmer et al., 2015), and *CD11c-Cre-IRF8^fl/fl^* (Sichien et al., 2016), IRF8-GFP(Wang et al., 2014), *Cish^−/−^*/IRF8-GFP. Mice with OVA specific TCR transgenic CD4^+^ T cells (OT-II) or CD8^+^ T cells (OT-I), GFP-OT-1 were housed under specific pathogen-free conditions at The Walter & Eliza Hall Institute of Medical Research. Chimera mice were Ly5.1/Ly5.2 F1 irradiated with 2 doses of 550 rads and reconstituted with 2×10^6^ BM cells from Ly5.1 and *Cish^−/−^* mice. All experiments were performed following relevant guidelines and regulations that were approved by the Walter & Eliza Hall Institute of Medical Research animal ethics committee (Project #2016.014, #2017.008, #2018.040).

### Cell Preparation, Antibodies, and Flow Cytometry

Cells from the spleen, lung and tumours were prepared by digestion in collagenase/DNase I. Antibodies (Abs) used in this study are listed in Supplementary Table 4. Cell numbers were determined by the addition of fluorochrome-conjugated calibration beads (BD Biosciences, San Jose, CA) directly to the samples. For evaluation of expression level, fluorescence minus one (FMO) control was included. Data were collected using FACS Verse (BD Biosciences) and analyzed using FlowJo software (Tree Star, Ashland, OR). Cell sorting was performed by using a FACS Aria or an Influx cell sorter (BD Biosciences).

### BM Cell Culture

BM cells from mice were isolated by flushing femurs and tibias with 5 ml PBS supplemented with 2% heat-inactivated fetal bovine serum (FBS) (Sigma Aldrich, Lenexa, KS, USA). The BM cells were centrifuged once and then re-suspended in tris-ammonium chloride at 37°C for 30 s to lyse red blood cell. The cells were centrifuged again and then strained through a 70-μm filter before being re-suspended in RPMI-1640 supplemented with 10% FBS. For GM-CSF stimulated culture, BM cells were re-suspended at 0.5 × 10^6^/ml containing titrated doses of GM-CSF in 12 well plates. After 3–4 days, the cultures were added fresh media with cytokines. Cell cultures were harvested on different day over 7 days. For further analysis, MØ and DCs were sorted on FACSaria (BD Biosciences).

### T cell Proliferation Assays

CD4^+^ T cells (OT-II) and CD8^+^ T cells (OT-I) were purified from TCR transgenic mice were purified by sorting and labeled with CTV (Invitrogen, ThermoFisher, Waltham, MA) (as per manufacturer's protocol). Labeled cells were cultured at 1 × 10^5^ in 200 μL RPMI1640 supplemented with 10% FBS in flat-bottom 96-well plates in the absence or the presence of antigen-presenting cells and defined antigens. Cell cultures were harvested 3 days later. T cell proliferation was evaluated by dye dilution. In some cultures, inhibitors of Arginase 1 L-Norvaline (12 mM, Sigma-Aldrich) was included. Cytokine levels in culture supernatants were analyzed using BioPlex kit (Biorad). In some experiments, CTV labeled CD8^+^ T cells were also stimulated with 5 μg/ml CD3 and 2 μg/ml anti-CD28 for 3 days. Cell proliferation and cytokine production were evaluated as above.

### Antigen uptake and processing

For uptake, one million of cells were incubated with 100 μg/mL FITC‐OVA at 37°C or kept on ice for 30 min. For processing, one million of cells were incubated with 100 μg/mL DQ-OVA (ThermoFisher) at 37°C for 30 min. Then samples were either incubated at 37°C for further 90 min or kept on ice. After incubation, cells were washed with cold 2% FCS‐EDTA (0.02 mM) PBS and analyzed by flow cytometry.

### Cell signaling analysis by Immunoblotting

GM-MØs were isolated by FACS sorting. The purity of enriched cells was consistent>95%. For cell signalling, cells were stimulated in vitro with recombinant GM-CSF (50 ng/ml) for indicated times in RPMI1640, supplemented with antibiotics and 10% heat-inactivated FCS. Cells were washed in cold PBS and pelleted followed by lysis. Approximately 5×10^5^ sorted cells were collected per sample and lysed in 100 μl lysis buffer (50mM Tris-HCl, pH 7.4, 150mM NaCl, 0.25% deoxycholic acid, 1% NP-40, 1mM EDTA) supplemented with protease inhibitors (Complete Cocktail tablets, Roche), 1 mM PMSF, 1 mM Na_3_VO_4_ and 1 mM NaF and incubated for 30 min on ice. Lysates were clarified by centrifugation at 13,000 rpm for 15 min at 4°C. Gel electrophoresis was carried out in 4-12% NuPAGE Bis-Tris gels followed by Western blotting to nitrocellulose membrane (Amersham). Primary antibodies are listed in Supplementary Table 4.

### Cell Stimulation and Cytokine Assay

MØs and moDCs derived from 7-day BM cultures were purified by flow sorting. Then cells were cultured at 5 × 10^4^ in 200 μL RPMI1640 supplemented with 10% FBS in U-bottom 96-well plates in the absence or the presence of LPS (1 μg/mL, Sigma) or CpG ODN 1668 (1 μM, Invivo Gen), agonistic anti-CD40 Ab (5 μg/ml, clone FGK4.5), or Poly I:C (5 μg/ml, InvivoGen) for 20 h. For cytokine detection from the supernatants of *in vitro* assays, the indicated cytokines were detected using BioPlex kit (Biorad). For intracellular cytokine staining, Cells were stimulated for 4–6 h with 0.5 μM CpG or 50 ng/ml PMA/1 mM ionomycin with GolgiStop (BD). Then, cells were surface stained for cell surface markers, fixed/ permeabilized using a Cytofix/Cytoperm kit (BD), and stained with anti-cytokine antibodies.

### Jak inhibition and IL-4 neutralization during in vitro MØ differentiation

For Jak inhibition, BM cells were cultured at 0.5 × 10^6^/ml containing 10 ng/mL GM-CSF in 12 well plates for 5 days. the cultures were added ruxolitnib (RUXO) (0.2 μM, from Invivo Gen). Cell cultures were harvested on 7 days for GM-MØs. For IL-4 neutralization, BM cells were cultured at 0.5 × 10^6^/ml containing 10 ng/mL GM-CSF in 12 well plates in the presence of anti-IL-4 Abs (50 μg/ml, clone 11B11) for 7 days.

### Tumour induction

1-5×10^5^ GM-CSF-transduced ouse melanoma B16F10 (B16-GM, provided by Prof. Jose Villadangos with permission from Prof Glenn Dranoff (Dranoff et al., 1993), were injected s.c into flank for 1-2 wks.

### RNA sequencing

GM-DCs and GM-MØs from 4 individual mice were enriched by sorting. RNA was isolated independently from biological replicates with RNeasyPlus Mini kits (Qiagen). mRNA reverse transcription and cDNA libraries were prepared using the TruSeq RNA Sample preparation kit (Illumina) following the manufacturer’s instructions. Samples were sequenced with an Illumina NextSeq 500 sequencing system, producing between 14-20 ×10^7^ single-end 85 bp reads per sample.

### Bioinformatic analysis of RNA-seq data

The RNA-seq reads were quantified against the Ensembl (Yates et al., 2020) v31 GRCm38 transcriptome using Kallisto (Bray et al., 2016). Differential expression analysis was conducted using Sleuth(Roguev et al., 2013) at the gene level via the method of Pimentel et al.(Yi et al., 2018). Q-values were calculated using Benjamini–Hochberg (Yoav Benjamini, 1995) adjustment, and log-fold changes at the gene level estimated by combining transcript level abundances normalized to Transcripts per Million reads (TPM) (Wagner et al., 2012). Heatmaps of the TPM expression values were plotted with the coolmap function from the limma package (Ritchie et al., 2015).

### Sample preparation for mass spectrometry-based proteomics

GM-MØs (1-1.5 million/replicate) from 4 individual WT or *Cish^−/−^* mice were sorted on a FACSaria to achieve a final purity of 99–100%. Cells were washed three times with ice cold PBS prior to dry cell pellet storage at − 80 °C. Cells were lysed in preheated (95°C) 5% SDS/10 mM tris/10 mM tris (2-carboxyethyl) phosphine/5.5 mM 2-chloroacetamide and heated at 95°C for 10 min. Neat trifluoracetic acid (Sigma) was added to hydrolyze the DNA, resulting in a final concentration of 1%. Lysates were quenched with 4 M tris (pH 10), resulting in a final concentration of ~140 mM tris (pH 7). Myeloid cell protein lysates (~50 μg) were prepared for mass spectrometry (MS) analysis as previously described (Rautela et al., 2019). Acidified peptide mixtures were analyzed by nanoflow reversed-phase liquid chromatography tandem mass spectrometry (LCMS/MS) on an Easy-nLC 1000 system (Thermo Fisher Scientific) coupled to a Q-Exactive HF (QE-HF) mass spectrometer equipped with a nanoelectrospray ion source and in-source column heater (Sonation) at 40 °C for automated MS/MS (Thermo Fisher Scientific). Peptide mixtures were loaded in buffer A (0.1% formic acid, 2% acetonitrile, Milli-Q water), and separated by reverse-phase chromatography using C_18_ fused silica column (packed emitter, internal diameter 75 μm, outer diameter 360 μm × 25 cm length, IonOpticks) using flow rates and data-dependent methods as previously described (Delconte et al., 2016) . Raw files consisting of high-resolution MS/MS spectra were processed with MaxQuant (version 1.5.8.30) for feature detection and protein identification using the Andromeda search engine as previously described (Rautela et al., 2019). LFQ quantification was selected, with a minimum ratio count of 2. PSM and protein identifications were filtered using a target-decoy approach at an FDR of 1%. Only unique and razor peptides were considered for quantification with intensity values present in at least 2 out of 3 replicates per group. Statistical analyses were performed using LFQAnalyst (https://bioinformatics.erc.monash.edu/apps/LFQ-Analyst/) whereby the LFQ intensity values were used for protein quantification. Missing values were replaced by values drawn from a normal distribution of 1.8 standard deviations and a width of 0.3 for each sample (Perseus-type). Protein-wise linear models combined with empirical Bayes statistics were used for differential expression analysis using Bioconductor package Limma whereby the adjusted *p*-value cutoff was set at 0.05 and log2 fold change cutoff set at 1. The Benjamini-Hochberg (BH) method of FDR correction was used.

### CRISPR/CAS9 mediated deletion of *Cish* in human CD34^+^ cord blood cells

CD34^+^ cells were isolated from human cord blood (Stemcell Technologies #17896) and cultured for 2 days in expansion media (Miltenyi #130-100-473). CIS RNPs were assembled by incubating 1ml of 30mM CIS sgRNA (5’-CTCACCAGATTCCCGAAGGT-3’; Synthego), 1.7ml of 67nM Cas9 (Integrated DNA Technologies #1081058), 1ml of electroporation enhancer (Integrated DNA Technologies #1075915) and 1.3ml PBS for 10min at room temperature. Five hundred thousand cells were pelleted and resuspended in RNP solution and 20ml of primary cell P3 buffer (Lonza), and were transferred to an electroporation curvette (Lonza). Cells were electroporated using the 4D-Nucleofector (Lonza) with pulse code CM-137 and rested in complete media for 10min before being transferred to cell culture dish. Cells were cultured with 5 ng/mL huGM-CSF (R&D) for 7 days. A small pellet of cells was taken for sequencing to determine CIS indel frequency.

### Cell transfer

WT and *Cish^−/−^* GM-MØs were incubated with 10 mg/ml OVA in serum free media at 37°C for 30 min. After wash with cold PBS, OVA-pulsed WT and *Cish^−/−^* GM-MØs were injected i.v into B6 mice or *Rag1−/−*/*Il2rg−/−* mice infused with 10^6^ Ly5.1-OT-1 2 weeks prior. OT-1 expansion was evaluated 3-7 days after GM-MØ injection.

### CUT&Tag-sequencing

The CUT&Tag was performed via Hyperactive in situ ChIP Library Prep Kit (Vazyme) following manufacturer recommendations. Briefly, *Cish^−/−^* GM-MØs were sorted from GM-CSF-supplemented cultures of BM cells from WT *IRF8^GFP^* and *Cish^−/−^ IRF8^GFP^*. GFP^+^ GM-MØs (1 × 10^5^ cells) were bound to concanavalin A cated magnetic beads (Bags laboratories) and were subjected for immunoprecipitation with 1ul (0.5μg) of primary antibody (rabbit anti-GFP, ab290, or rabbit anti-H3k27me3, CST:9733, or rabbit anti-mouse IgG control). Following primary incubation and washing using a magnetic stand, a secondary anti rabbit antibody was added and incubated under gentle agitation for 1h (Antibodies online ABIN101961). Cells were washed and incubated for 1h in a mix of Hyperactive pG-Tn5/pA-Tn5 Transposon with Dig-300 Buffer to a final concentration of 0.04 μM. Excessive reagents were washed using a magnetic stand, and cells were resuspended in 100ul of Tagmentation buffer and incubated at 37°C for 1h. Reaction was stopped by heat inactivation (55°C for 10 min). DNA was purified using Ampure XP beads (Beckman coulter). For library amplification, 24ul of DNA was mixed with 10ul of 5 x TAB, 1ul of TAE, 5ul of uniquely barcoded i5 and i7 primers (Mezger et al., 2018), and amplified for 14 cycles (72 °C for 5 min; 98 °C for 30 s; 14 cycles of 98 °C for 10 s, 63 °C for 10 s and 72 °C for 1 min; and hold at 4 °C). PCR products were purified with Ampure XP beads and eluted in 25ul water. Eluted DNA was checked for region molarity via High sensitive D5000 tap station (Sup Fig 6B). Libraries were sequenced on an Illumina NextSeq platform and 150-bp paired-end reads were generated.

### Cut&Tag data analysis

The raw data was uploaded on Galaxy Australia website (https://usegalaxy.org.au/). The sequence quality was checked via FastQc and MultiQc. Then the raw data were aligned to mm10 via Bowtie2 version 2.3.4.3 with options: --local --very-sensitive-local --no-unal --no-mixed --no-discordant -- phred33 −I 10 −X 700 (Langmead and Salzberg, 2012; Langmead et al., 2009). Duplicate reads were removed via SAM/BAM filter. The biological duplicate sample reads were pooled together via Merge BAM files package. Then peaks were called via MACS2 (q<0.01)(Feng et al., 2012; Zhang et al., 2008). The binding peaks were viewed using the Integrated Genome Brower (IGB)(Nicol et al., 2009). The peaks were annotated tvia ChIPseeker (Yu et al., 2015). The peak intersection between WT and KO macrophages was calculated via Bedtool (Quinlan and Hall, 2010).

### Retroviral transduction

Retroviral supernatants were generated by transient transfection of 293T cells with plasmids encoding viral envelope proteins (pMD1-gag-pol and pCAG-Eco), and expression vectors encoding for pMSCV-iresGFP and pMSCV-IRF8iresGFP using FuGeneHD (Promega). Retroviral supernatants were centrifuged onto RetroNectin (Takara)-coated plates for 45 min at 4000 rpm at 32°C. Cells were then cultivated with the virus in the presence of 4 μg/ml polybrene (Sigma-Aldrich) for 12 hr.

### Cell imaging

The 2×10^5^/ml cells were suspended in 100 μl PBS and stained using 1:1000 dye mix (CellTracker velvet, LysoTracker green and DAPI, from Thermo Fisher) at 37 ℃ for 15 min, then washed twice in PBS and placed in 96 well PerkinElmer Cell Carrier plates for imaging. A Leica SP8 confocal microscope was used. The volume of each cell was calculated using Imaris 9.1.2 by applying a surface to each cell using the CellTracker velvet channel. Images of higher resolution were taken using the same microscopy setting and processed on FIJI.

### Allergic asthma model

To build allergic asthma, WT or *Cish^−/−^* GM-MØs were loaded with OVA, then were injected into the recipient mice at day 0 and day7. The allergic response were induced as described previously(Keenan et al., 2019).

### Statistical Analysis

Statistical comparisons of mean difference between two groups from independent experiments were made using a student's *t*-test, and data presented as multiple groups, dose-response curves or time courses were analyzed using ANOVA. The analysis was performed with Prism v.5.0 software (GraphPad, San Diego, CA). P< 0.05 were considered statistically significant.

### Data Availability

The RNA-seq data have been deposited to the European Nucleotide Archive (ENA) with dataset identifier PRJEB40745, while the mass spectrometry proteomics data have been deposited to the ProteomeXchange Consortium via the PRIDE partner repository with the dataset identifier PXD018390 (reviewer token: Username: Password:2MzhUnon). CUT&Tag data will be available upon request and will be publicly available after publication.

## Acknowledgments

We thank Lisa Reid, Marina Patsis, Manuela Hancock, Rhiannan Crawley, Rebekah Meeny, Tania Camilleri and the institute flow cytometry facility for excellent technical assistance. We acknowledge the Wurundjeri people of the Kulin nation as the traditional owners and custodians of the land on which most of the work was performed. This work was supported by National Health and Medical Research Council of Australia (NHMRC) grants (1037321, 1105209, 1143976, 1150425, 1080321, 5575500, 1054925, 1048278), NHMRC Independent Research Institutes Infrastructure Support Scheme grant (361646) and Victorian State Government Operational Infrastructure Support grant. J.B. was supported by the Stafford Fox Medical Research Foundation.

## Authors contributions

Conceptualization, YZ; Methodology: SBZ, JR, MC, NGB, LFD, JB, YY,YX; Investigation: YZ, SBZ, TBK, JR, MC, QKW, HQW, LS, RS, LFD, FSFG, YY, YX, RA, NI; Writing – Original Draft, YZ with input from SBZ,AML, SN, SEN, MC, NGB, and YY; Review & Editing: SBZ, MC, NGB, LFD, YKX, SN, NDH, SEN, AML; Resources: JR, NDH, SN, SEN.

