## Supplemental figures and tables for "CIS calibrates GM-CSF signalling strength to regulate macrophage polarization via a STAT5-IRF8 axis": Supplementary Figures CIS manuscript Zhang S et al.pdf

### Fig S1, related to Figure 1

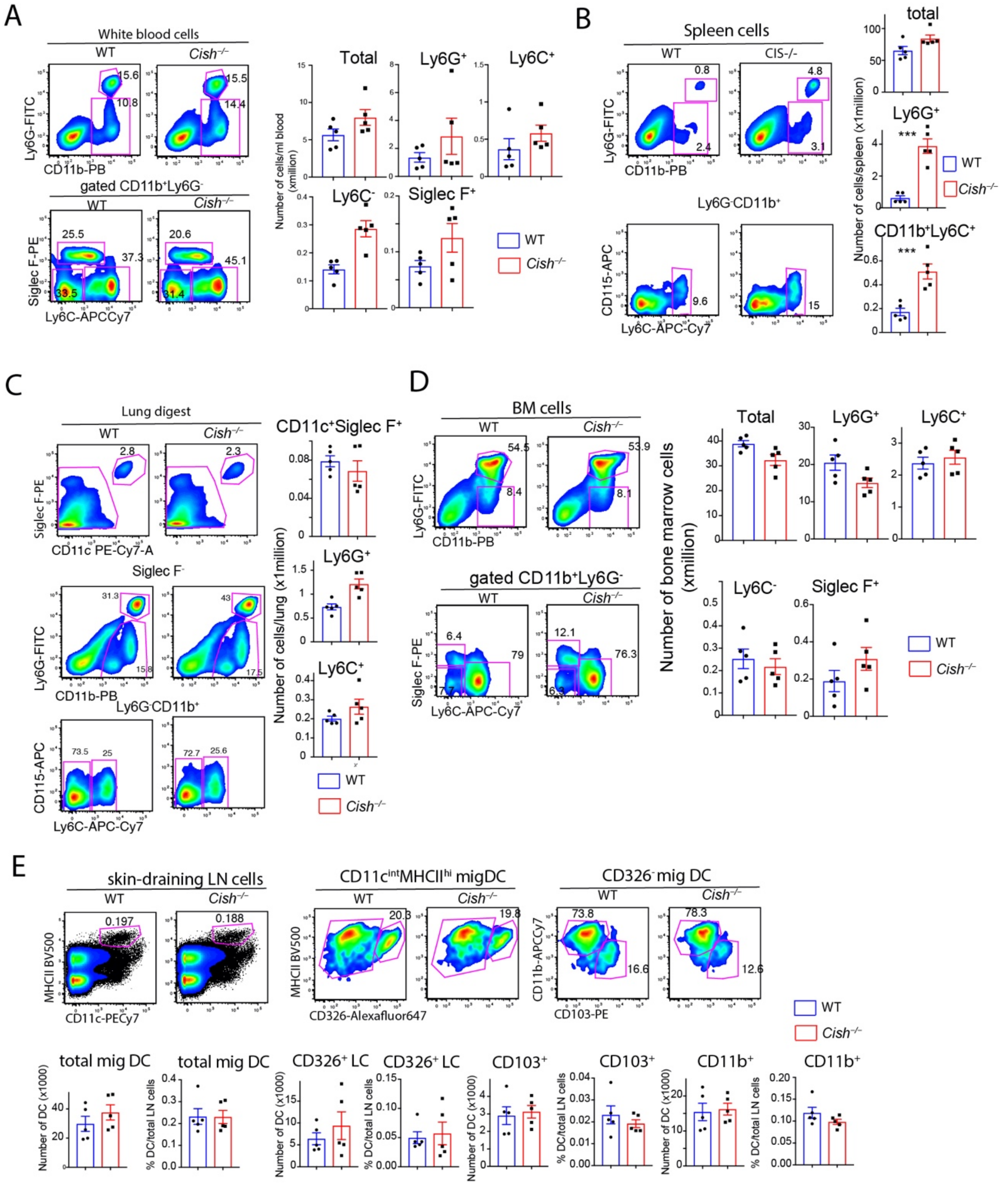


**Fig S1. *Cish*$^{-/-}$ mice has normal distribution of multiple types of myeloid cells**. Blood, spleen, lung, bone marrow and skin-draining lymph node from C57BL/6 (n=5) and *Cish*$^{-/-}$ mice (n=5) were analysed for myeloid cells. **(A)** Neutrophils (Ly6G$^{+}$), Monocytes (Ly6C$^{+}$ or Ly6C$^{-}$) and Eosinophils (SiglecF$^{+}$) in blood. **(B)** Ly6C$^{+}$ Monocytes and Neutrophils in spleen, **(C)** Alveolar macrophages (CD11c$^{+}$SiglecF$^{+}$), Ly6C$^{+}$ Monocytes and Neutrophils in lung. **(D)**

Neutrophils, Monocytes (Ly6C$^+$ or Ly6C$^-$) and Eosinophils in bone marrow. **(E)** MHCII$^{hi}$CD11c$^+$ migratory DCs and three subsets (CD326$^+$, CD103$^+$ and CD11b$^+$) in lymph node. Bar graphs show mean+/-SEM of indicated populations. **P<0.01, ***P<0.001 by t-test.

Figure S2, related to Figure 1

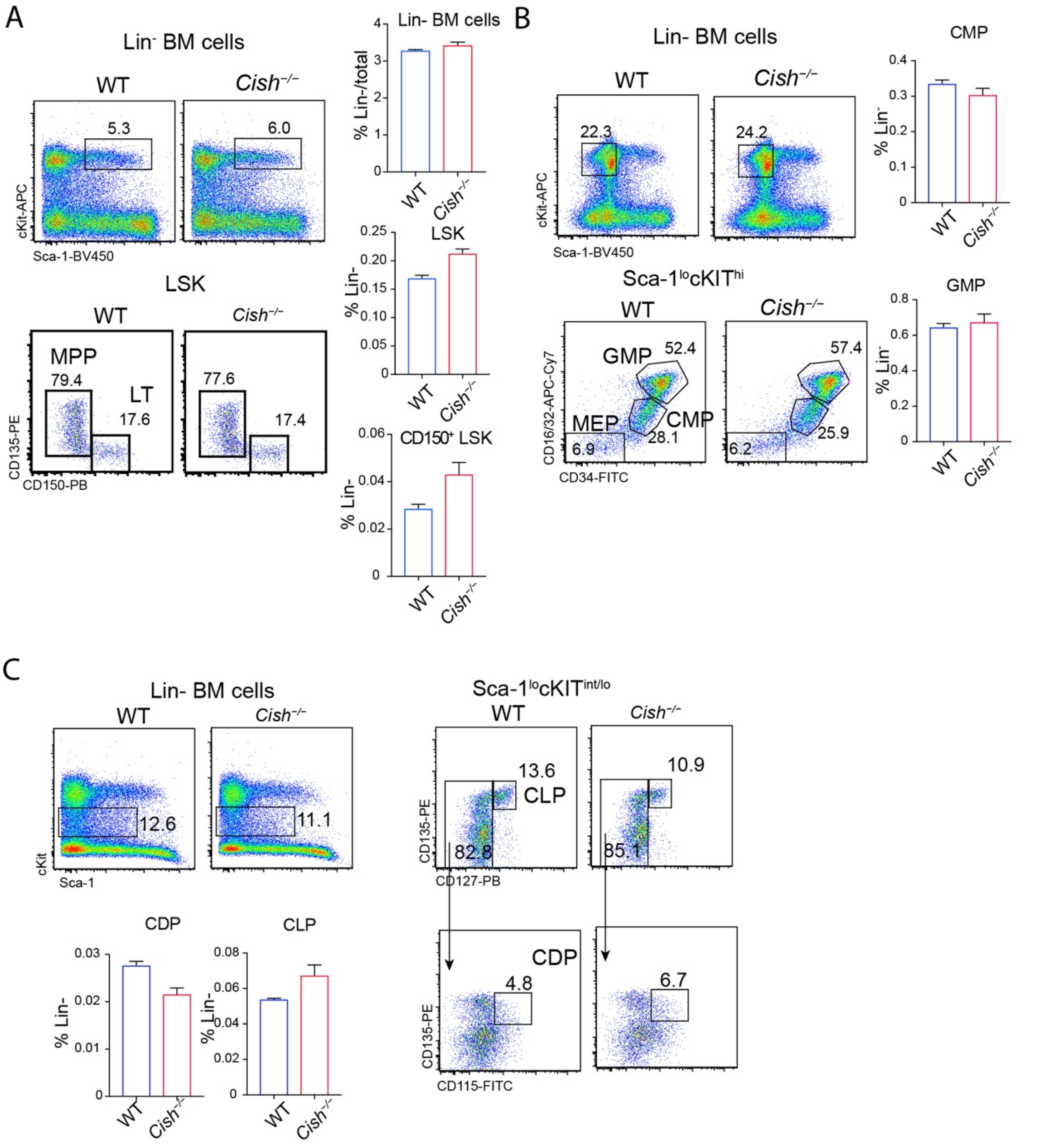


**Fig S2. *Cish*$^{-/-}$ mice has normal distribution of BM hematopoietic progenitors**. BM cells from individual mice (n=4) were analyzed for BM hematopoietic progenitors. **(A)** Lin-Sca1$^{hi}$cKIT$^{hi}$ cells (LSK) cells are shown for multiple progenitors (MPP, CD150$^{+}$CD135$^{-}$) and long-term HSCs (CD150+CD135-). **(B)** Lin-Sca1$^{lo}$cKIT$^{hi}$ cells are shown for GMP (CD16/32$^{hi}$CD34$^{hi}$), CMP (CD16/32$^{int}$CD34$^{int}$), and MEP (CD16/32$^{lo}$CD34$^{lo}$). **(C)** Lin-Sca1$^{lo}$cKIT$^{int}$ cells are shown for CLP (CD135$^{+}$CD127$^{+}$) and CDP (CD135$^{+}$CD127$^{-}$

CD115$^{+}$). **(D)** Bar graphs show mean+/-SEM of numbers of indicated populations. Data from one of two experiments are shown.

### Figure S3, related to Figure 1

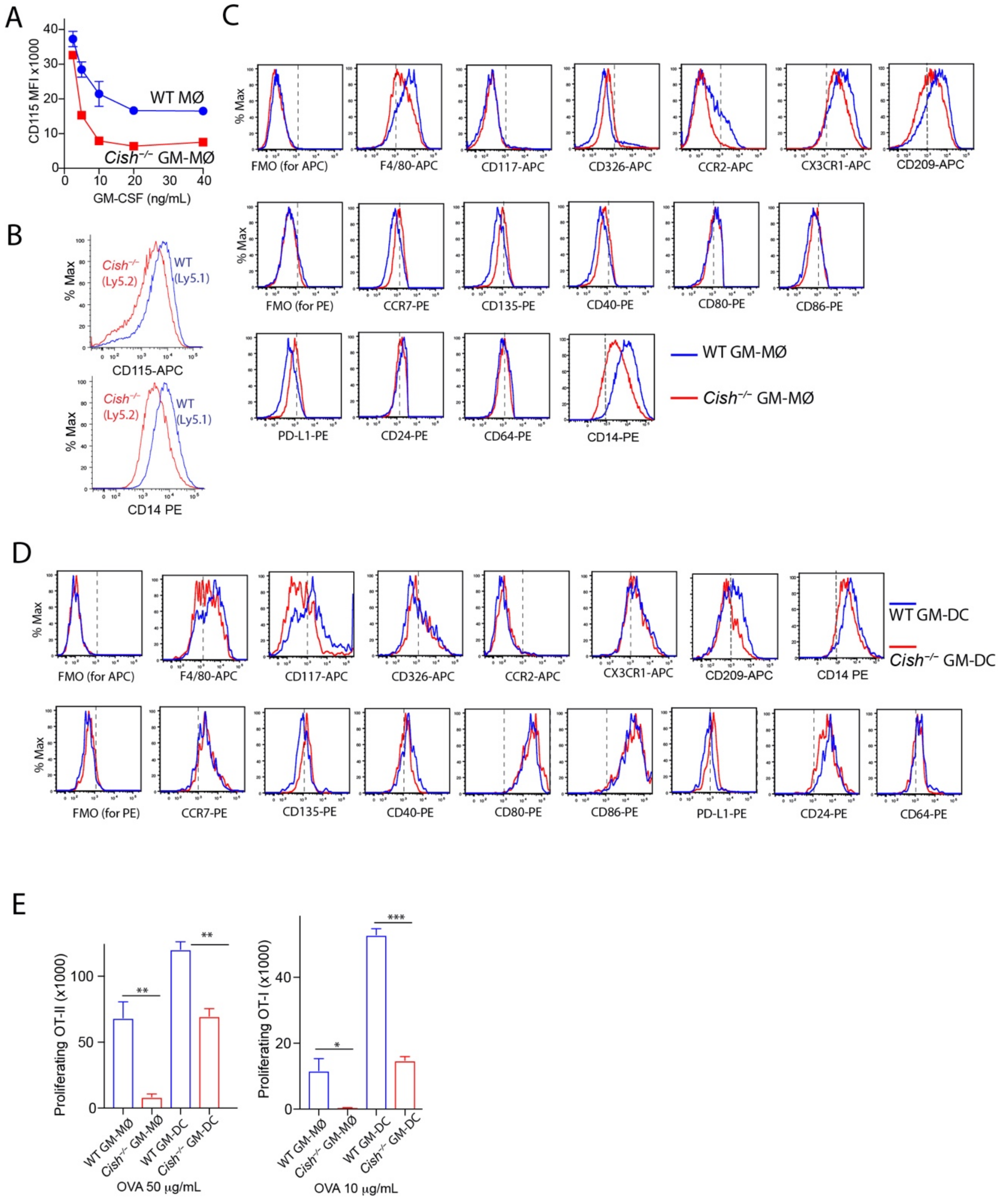


**Fig. S3. Cish deficiency alters the expression of surface markers and function by GM-MØs and GM-DCs.** (**A**) dose-dependent downregulation of CD115 by *Cish*$^{-/-}$ GM-MØs. GM-MØs derived with different GM-CSF concentrations were analyzed for CD115 expression. Line plot shows MFI of CD115 of WT and *Cish*$^{-/-}$ GM-MØs. (**B**) Cell-intrinsic downregulation of CD115 and CD14 by *Cish*$^{-/-}$ GM-MØs. BM cells from WT (Ly5.1) and *Cish*$^{-/-}$ (Ly5.2) mice were co-cultured with 10 ng/mL GM-CSF for 7 days. Gated GM-MØs are shown for expression of CD115 and CD14. (**C**) Expression of the indicated myeloid markers in WT and *Cish*$^{-/-}$ GM-MØs . (**D**) Expression of the indicated myeloid markers in WT and *Cish*$^{-/-}$ GM-DCs. (**E**) Reduced T-cell stimulation by both GM-MØs and GM-DCs with CIS deficiency. Purified GM-MØs and GM-DCs of WT and *Cish-/-* origin were co-cultured with CTV-labelled OT-II CD4$^{+}$ T cells for 3 days or OT-1 CD8$^{+}$ T cells for 2 days in the presence or absence of indicated concentrations of OVA proteins *P<0.05, **P<0.01, ***P<0.001 by multiple group comparison of ANOVA test.

### Figure S4, related to Figure 2

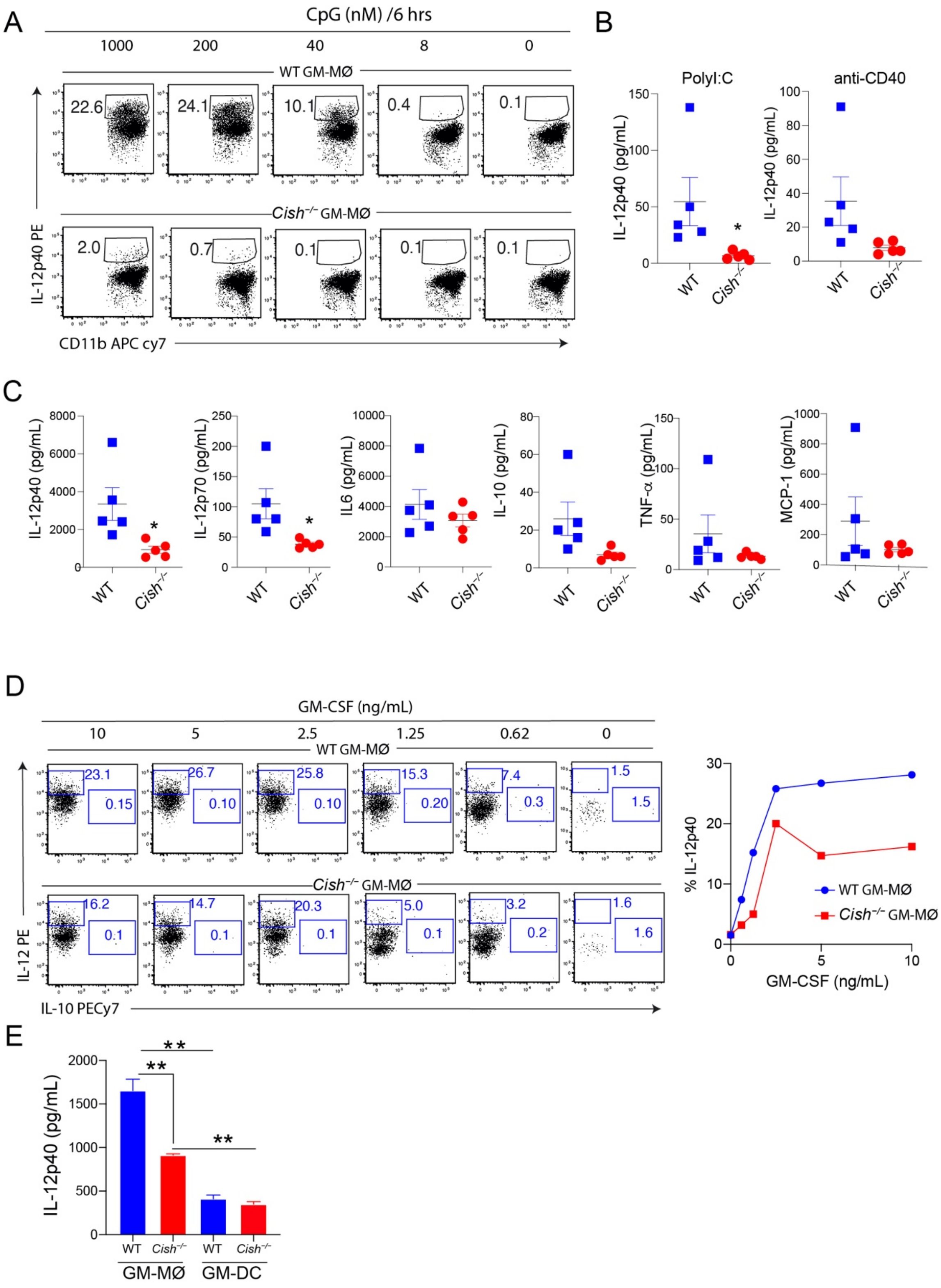


**Fig. S4.** ***Cish*$^{-/-}$ MØs are defective in IL-12 production.** (**A**) IL-12p40 production by WT and *Cish*$^{-/-}$ GM-MØs stimulated with different concentration of CpG. (**B**) IL-12p40 production by WT and *Cish*$^{-/-}$ GM-MØs derived from individual mice were stimulated by PolyI:C or agonist anti-CD40 Ab. Means and SD of IL-12p40 are shown. (**C**) Production of IL-12p40/p70, IL-6, IL-10, TNF-α and MCP-1 under CpG stimulation by WT and *Cish*$^{-/-}$ GM-MØs derived from individual mice. (**D**) IL-12p40 production evaluated by intracellular cytokine staining after CpG stimulaton for 4 hrs by WT and *Cish*$^{-/-}$ GM-MØs derived with different concentrations of GM-CSF. (**E**) IL-12p40 production by GM-MØs and GM-DCs from WT and *Cish*$^{-/-}$ cultures were evaluated after CpG stimulation for 20 hrs. Means and SD of IL-12p40 are shown. **P<0.01 between indicated comparisons by multiple comparison of ANOVA test.

### Figure S5, related to Figure 3

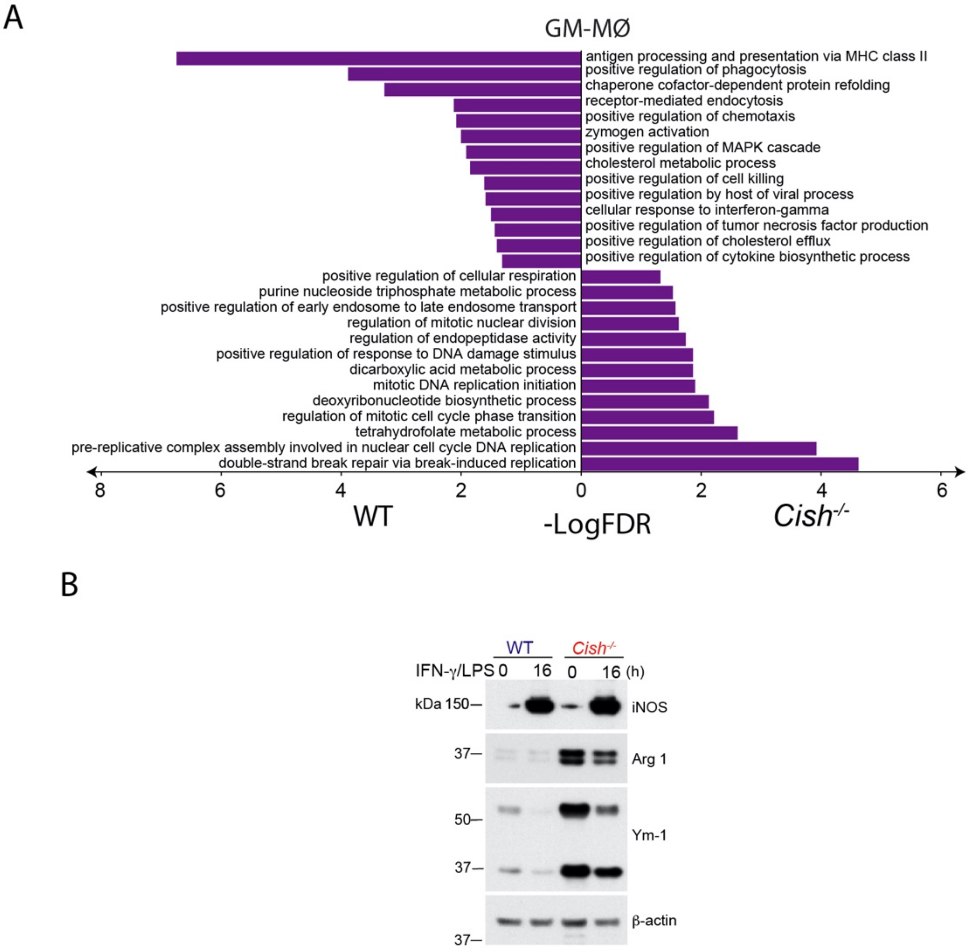


**Fig. S5. *Cish*$^{-/-}$ MØs have characteristics of M2 MØs.** (**A**) GO enrichment analysis of biological processes for up and down-regulated genes between WT and *Cish*$^{-/-}$ GM-MØs. (**B**) WT and *Cish*$^{-/-}$ GM-MØs were stimulated with IFN-γ plus LPS for 16 hrs. Harvested WT and *Cish*$^{-/-}$ GM-MØs were then analysed for Ym1 and Arg1 expression by Western Blot.

Supplemental Figure 6 (related to Figure 6)

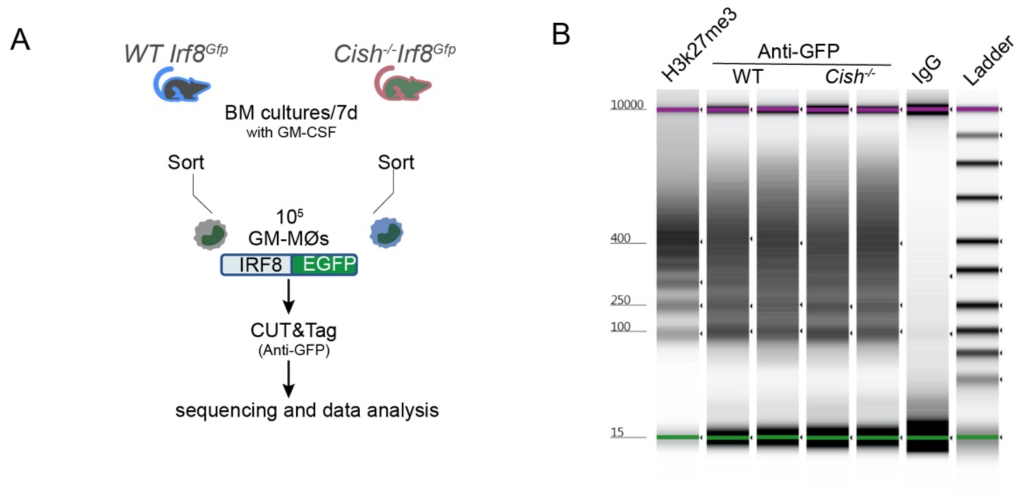


**Fig. S6. IRF8 target genes. (A)** Flow chart for CUT&Tag to analyze IRF8 binding peaks in GM-MØs. (**B**) Size distribution of fragment of the DNA library via the tape station high sensitive D5000. Two biological repeats for both WT and *Cish*$^{-/-}$ GM-MØs were included. The H3k27me3 was used as positive control and the IgG was used as negative control.

**Supplementary Table 4**

**Antibodies, biological materials and resources**

| REAGENT or RESOURCE | SOURCE | IDENTIFIER |
|---|---|---|
| **Antibodies (Clone)** | | |
| Anti-mouse CD326-Alexafluor647 (G8.8) | BioLegend | Cat#118212, *RRID*:AB_1134101 |
| Anti-mouse CD103-PE (M290) | BD | Cat#557495, *RRID*:AB_396732 |
| Anti-mouse CD117-APC (2B8) | BD | Cat#553356, *RRID*:AB_398536 |
| Anti-mouse Ly6c-APCCy7 (AL-21) | BD | Cat#560596, *RRID*:AB_172755 |
| Anti-mouse CD11b-PB (M1/70) | In house | N/A |
| Anti-mouse Ly6G-FITC (1A8) | In house | N/A |
| Anti-mouse CD115-APC (T38-320) | BD | Cat#567027, *RRID*:AB_2870013 |
| Anti-mouse Siglec F-PE (E50-2440) | BD | Cat#552126, *RRID*:AB_394341 |
| Anti-mouse CD4-PE-Cy7 (RM4-5) | BD | Cat#552775, *RRID*:AB_394461 |
| Anti-mouse CD4-BV500 (RM4-5) | BD | Cat#560782, *RRID*:AB_1937315 |
| Anti-mouse CD206-APC (C068C2) | Biolegend | Cat#141707, *RRID*:AB_10896057 |
| Anti-mouse Sca1-PacificBlue (E13-161.7) | Biolegend | Cat#122520, *RRID*:AB_2143237 |
| Anti-mouse CD150-PB (TC15-12F12.2) | Biolegend | Cat#115924, *RRID*:AB_2270307 |
| Anti-mouse CD135-PE (A2F10) | eBioscience | Cat#12-1351, *RRID*:AB_465859 |

| | | |
|---|---|---|
| Anti-mouse CD34-FITC (RAM34) | BD | Cat#560238, *RRID*:AB_1645242 |
| Anti-mouse CD16/32-APCCy7 (2.4G2) | BD | Cat#560541, *RRID*:AB_1645229 |
| Anti-mouse CD127-PB (A7R34) | eBioscience | Cat#48-1271,*RRID*:AB_2016629 |
| Anti-mouse CD115-PE (T38-320) | BD | Cat#565249, *RRID*:AB_2739132 |
| Anti-mouse F4/80-APC (BM8) | Bioscience | Cat#17-4801, *RRID*:AB_469452 |
| Anti-mouse CCR2-APC (475301) | R&D systems | Cat#FAB5538A,*RRID*:AB_10645617 |
| Anti-mouse CCR2-PE (475301) | R&D systems | Cat#FAB5538P,*RRID*:AB_10718414 |
| Anti-mouse CX3CR1-APC (SAO11F11) | BioLegend | Cat#149008, *RRID*: AB_2564491 |
| Anti-mouse CD209-eFluor660 (MMD3) | eBioscience | Cat#50-2094-80, *RRID*: AB_11219065 |
| Anti-mouse CD86-PE (GL1) | BD | Cat#553692, *RRID*:AB_394994 |
| Anti-mouse CD80-PE (16 10A1) | BD | Cat#553691, *RRID*:AB_394993 |
| Anti-mouse PD-L1-PE (MIH5) | BD | Cat#558091, *RRID*:AB_397018 |
| Anti-mouse CD24-PE (M1/69) | BD | Cat#*553262*, *RRID*:AB_394741 |
| Anti-mouse CD64-PE (X54-5/7.1) | BioLegend | Cat#139304, *RRID*:AB_10612740 |
| Anti-mouse CD14-PE (rmC5-3) | BD | Cat#553740, *RRID*:AB_395022 |

| | | |
|---|---|---|
| Anti-mouse TCRVa2-PE (B20.1) | BD | Cat#553289, *RRID*:AB_394760 |
| Anti-mouse I-A/I-E-BV500 (M5/114.15.2) | BD | Cat#562366, *RRID*:AB_11153488 |
| Anti-mouse CD11c-PECy7 (HL3) | BD | Cat#558079, *RRID*:AB_647251 |
| Anti-mouse CD8-APCCy7 (53-6.7) | BD | Cat#557654, *RRID*:AB_396769 |
| Anti-mouse IFNg-APC-eFluor780(XMG1.2) | eBioscience | Cat#47-7311-82, *RRID*:AB_2688061 |
| Anti-mouse IFNg-FITC (XMG1.2) | eBioscience | Cat#11-7311, *RRID*:AB_465412 |
| Anti-mouse Siglec H-PE (eBio440c) | eBioscience | Cat#12-0333, *RRID*:AB_10597139 |
| Anti-mouse mouse Fc R1a-PECy7 (1-Mar) | eBioscience | Cat#25-5898, *RRID*:AB_2573493 |
| Anti-mouse GM-CSFR-APC | In house | N/A |
| Anti-mouse GM-CSFRα-APC (698423) | R&D systems | Cat#FAB6130A, *RRID*:AB_10973836 |
| Rat anti-TNP-KLH-PE (A95-1) | BD | Cat#553989, *RRID*:AB_10049479 |
| Rat anti-TNP-KLH-APC (A95-1) | BD | Cat#556924, *RRID*:AB_10063104 |
| Anti-mouse XCR1-BV670 (ZET) | BioLegend | Cat#148220, *RRID*:AB_2566410 |
| Anti-mouse IL12p40-Biotin (C17.8) | BD | Cat#554476, *RRID*:AB_395419 |
| Strept-PerCP | BD | Cat#554064, *RRID*:AB_2336918 |

| | | |
|---|---|---|
| Anti-mouse CD11b-APCCy7 (M1/70) | BD | Cat#557657, *RRID*:AB_396772 |
| Anti-mouse IL12p40-PE (C17.8) | BD | Cat#562038,<br>*RRID*:AB_10895571 |
| Anti-mouse IL10-PECy7 (JES5-16E3) | BD | Cat#***505026,***<br>*RRID*:AB_11150582 |
| Anti-mouse CD45.1-FITC (A20) | BD | Cat#553775, *RRID*:AB_395043 |
| Anti-mouse CD45.2-FITC (104) | BD | Cat#553772, *RRID*:AB_395041 |
| Anti-mouse CD45.1-Pacific Blue (A20) | In house | N/A |
| Anti-mouse CD45.2 (104) | BioLegend | Cat#109820, *RRID*:AB_492872 |
| Anti-human HLA-DR-APCCy7 (L243) | BD | Cat#335796, *RRID*:AB_399974 |
| Anti-human CD14-FITC (M5E2) | BD | Cat#555397, *RRID*:AB_395798 |
| Anti-human CD16-PE (3G8) | BD | Cat#555407, *RRID*:AB_395807 |
| Anti-Arg1 | BD | Cat#610708, RRID:AB_398031 |
| Anti-Ym1 | STEM CELL | Cat#01404, RRID:AB_528565 |
| Anti-b-actin-HRP | Santa Cruz | Cat#sc-47778 |
| Anti-ERK1/2 | Cell Signaling | Cat#9102, RRID:AB_330744 |
| Anti-pERK1/2 | Cell Signaling | Cat#9101, RRID:AB_331646 |
| Anti-Akt1 | Cell Signaling | Cat#4691, RRID:AB_915783 |
| Anti-pAkt1 | Cell Signaling | Cat#9275, RRID:AB_329828 |
| Anti-CIS (D4D9) | Cell Signaling | Cat#8731 |
| Anti-STAT5 | Cell Signaling | Cat#94206 |
| Anti-pSTAT5 (8-5-2) | Millipore | Cat#05-495, RRID:AB_309762 |
| INOS | BD | Cat#61032 |
| **Bacterial and virus strains** | | |

| | | |
|---|---|---|
| NEB® 5-alpha Competent *E. coli* (High Efficiency) | New England Biolabs | Cat# C2987 |
| **Oligonucleotides** | | |
| PCR primers: mCIS F ex2: gacgttctcctaccttcgggaa | N/A | N/A |
| PCR primers: mCIS R ex3 aggaatgtaccctccggcatct | N/A | N/A |
| **Biological Samples** | | |
| CIS RNP guide 57 5'-CTCACCAGATTCCCGAAGGT-3' | Synthego | N/A |
| Cas9 | Integrated DNA Technologies | Cat#1081058 |
| electroporation enhancer | Integrated DNA Technologies | Cat#1075915 |
| 4D-Nucleofector | Lonza | AAF-1002B |
| **Chemicals, Peptides, and Recombinant Proteins** | | |
| Albumin from chicken egg white (OVA) | Sigma-Aldrich | Cat#A5503 |
| SIINFEKL | GL Biochem | Cat#SL-8; 53698 |
| OVA (323–339) | InvivoGen | Cat# vac-isq |
| Mouse GM-CSF | Peprotech | Cat#315-03 |
| Human GM-CSF | R&D | Cat#215-GM-500 |
| Ruxolitnib | Sigma-Aldrich | Cat#ADV390218177 |
| Collagenase | Worthington | Cat#CLS-3 |
| DNase | Sigma-Aldrich | Cat#10104159001 |

| | | |
|---|---|---|
| Cytofix/Cytoperm kit | BD Bioscience | Cat#554714 |
| Protease inhibitors | Roche | Cat#11697498001 |
| CpG ODN 1668 | Invitrogen | Cat# tlrl-1668 |
| LPS | Sigma-Aldrich | Cat# L8274 |
| L-Norvaline | Sigma-Aldrich | Cat# N7627-1G |
| Anti-murine GM-CSF (MP-1-22E9) | In house | N/A |
| DQ-OVA | Thermofischer | Cat#D12053 |
| FITC-conjugated OVA | In house | N/A |
| Anti-PE microbeads | Miltenyi Biotec | Cat#130-048-801 |
| Anti-FITC microbeads | Miltenyi Biotec | Cat#130-048-701 |
| **Critical Commercial Assays** | | |
| Cell Trace Violet proliferation kit | Thermo Fisher | Cat#C34557 |
| Mouse cytokine 23-plex Assay | Bio-rad | Cat#M60009RDPD |
| Human cytokine Screening Panel | Bio-rad | Cat#12007283 |
| RNeasyPlus Mini kits | Qiangen | Cat#74134 |
| TruSeq RNA Sample preparation kit | Illumina | Cat# RS122-2001 |
| Bio-plex pro mouse cytokine chemokine and growth factor assays 23-Plex panel | Bio-rad | Cat#M60009RDPD |
| BioMag Plus Concanavalin A | Bangs laboratories, Inc | Cat#BP531 |
| Hyperactive In-Situ ChIP library Prep Kit for Illumina | Vazyme | Cat#TD902 |
| AMPure XP beads | BECKMAN | Cat#A63880 |
| FugeneHD Transfection Reagent | Promega | Cat# E2312 |

| | | |
|---|---|---|
| **Deposited data** | | |
| Data will be made publicly accessible upon publication | | |
| **Experimental Models: Cell Lines** | | |
| B16-F10 | ATCC | CRL-6475 |
| B16-GM-CSF | (Dranoff et al., 1993) | N/A |
| **Experimental Models: Organisms/Strains** | | |
| Mouse: CD11c-Cre/Irf8$^{fl/fl}$ | (Sichien et al., 2016) | N/A |
| Mouse: IRF8-GFP reporter mice | (Wang et al., 2014) | N/A |
| Mouse: Tg(TcraTcrb)1100Mjb. (OT-I) | (Barnden et al., 1998) | N/A |
| Mouse: Tg(TcraTcrb)425Cbn (OT-II) | (Hogquist et al., 1994) | N/A |
| Mouse: GFP-OT-I | (Hogquist et al., 1994) | N/A |
| Mouse: IRF8-GFP/Cish$^{-/-}$ | This paper | N/A |
| Mouse: Rag1$^{-/-}$/Il2rg$^{-/-}$, | Internal | N/A |
| Mouse: Cish$^{-/-}$ | (Palmer et al., 2015) | N/A |
| Mouse: Ly5.1/J | Internal | N/A |
| Mouse: Ly5.1/Ly5.2 F1 | Internal | N/A |

| | | |
|---|---|---|
| Mouse: C57BL/6 | Internal | N/A |
| **Recombinant DNA** | | |
| MSCV IRES GFP | Addgene | Cat#20672 |
| MSCV IRF8 IRES GFP | (Chopin et al., 2019) | N/A |
| **Software and Algorithms** | | |
| Flowjo V9, V10 | FlowJo, LLC | https://www.flowjo.com/ |
| Prism 7.0, 8.0 | GraphPad Software | https://www.graphpad.com/scientific-software/prism/ |
| Integrated Genome Browser (IGB) | N/A | https://bioviz.org |
| Adobe Illustrature CC 2019 | Adobe | https://www.adobe.com/sea/products/illustrator.html?promoid=PGRQQLFS&mv=other |
| LFQAnalyst | N/A | https://bioinformatics.erc.monash.edu/apps/LFQ-Analyst/ |
| **Other** | | |
| BD FACSverse | BD Bioscience | N/A |
| BD Fortessa | BD Bioscience | N/A |
| BD FACS ARIA | BD Bioscience | N/A |
| Bio-plex 200 Sytems | Bio-Rad | 171000205 |
